# Generation of Schwann cell derived melanocytes from hPSCs identifies pro-metastatic factors in melanoma

**DOI:** 10.1101/2023.03.06.531220

**Authors:** Ryan M. Samuel, Albertas Navickas, Ashley Maynard, Eliza A. Gaylord, Kristle Garcia, Samyukta Bhat, Homa Majd, Mikayla N. Richter, Nicholas Elder, Daniel Le, Phi Nguyen, Bradley Shibata, Marta Losa Llabata, Licia Selleri, Diana J. Laird, Spyros Darmanis, Hani Goodarzi, Faranak Fattahi

## Abstract

The neural crest (NC) is highly multipotent and generates diverse lineages in the developing embryo. However, spatiotemporally distinct NC populations display differences in fate potential, such as increased gliogenic and parasympathetic potential from later migrating, nerve-associated Schwann cell precursors (SCPs). Interestingly, while melanogenic potential is shared by both early migrating NC and SCPs, differences in melanocyte identity resulting from differentiation through these temporally distinct progenitors have not been determined. Here, we leverage a human pluripotent stem cell (hPSC) model of NC temporal patterning to comprehensively characterize human NC heterogeneity, fate bias, and lineage development. We captured the transition of NC differentiation between temporally and transcriptionally distinct melanogenic progenitors and identified modules of candidate transcription factor and signaling activity associated with this transition. For the first time, we established a protocol for the directed differentiation of melanocytes from hPSCs through a SCP intermediate, termed trajectory 2 (T2) melanocytes. Leveraging an existing protocol for differentiating early NC-derived melanocytes, termed trajectory 1 (T1), we performed the first comprehensive comparison of transcriptional and functional differences between these distinct melanocyte populations, revealing differences in pigmentation and unique expression of transcription factors, ligands, receptors and surface markers. We found a significant link between the T2 melanocyte transcriptional signature and decreased survival in melanoma patients in the cancer genome atlas (TCGA). We performed an *in vivo* CRISPRi screen of T1 and T2 melanocyte signature genes in a human melanoma cell line and discovered several T2-specific markers that promote lung metastasis in mice. We further demonstrated that one of these factors, SNRPB, regulates the splicing of transcripts involved in metastasis relevant functions such as migration, cell adhesion and proliferation. Overall, this study identifies distinct developmental trajectories as a source of diversity in melanocytes and implicates the unique molecular signature of SCP-derived melanocytes in metastatic melanoma.

## Introduction

The neural crest (NC) is a transient, multipotent, fetal cell population specified along the dorsal neural tube that gives rise to diverse cell types, such as peripheral neurons and glia, melanocytes, cranial osteoclasts and chondrocytes^1^. During normal development, diversification of potency throughout the NC population remains a topic of extensive research^2^. Lineage tracing, chimeric grafts and explant experiments have provided compelling evidence that NC fate potential is both regionally and temporally restricted during normal development. The combination of a NC cell’s (NCC) position along the rostrocaudal axis and duration of migration determines its lineage options^3–9^. This points to cell-intrinsic differences in spatiotemporally distinct NCCs^10–12^.

Understanding the molecular programs underlying NC lineage potential and restriction has implications in basic stem cell and developmental biology as well as for the development of therapies for numerous neurocristopathies. However, the transient and migratory nature of NCCs pose challenges in tissue accessibility and scalability necessary for -omics level studies and high-throughput molecular perturbations, especially from human tissue. Established strategies for differentiating NCCs from human pluripotent stem cell (hPSC) have provided an alternative model system with the ability to pattern into cranial, vagal, trunk and sacral regional identities^13–15^. Recently, Majd et al. demonstrated temporal patterning of hPSC-NCCs with prolonged culture of 3D spheroids resulting in more efficient differentiation to glial lineages, mimicking the *in vivo* shift to gliogenesis in late-migrating NCCs^16^. This differentiation system enables the study of temporal fate restriction in NCCs, and promises access to lineages that emerge from late-migrating NCCs, such as parasympathetic neurons^17,18^. Interestingly, the melanocyte lineage is uniquely accessible to both early and late migratory NCCs^19,20^. However, further characterizations are needed to determine how temporally distinct differentiation trajectories affect melanocyte identity and function.

In this study, we characterized the temporal changes in hPSC-NCC populations using single cell transcriptomics, revealing changes in glial propensity and temporarily distinct melanogenic progenitors with unique transcription factor and signaling activity. Building on the protocol described by Majd et al., we developed a strategy for the directed differentiation of melanocytes from hPSCs through a Schwann cell precursor (SCP) intermediate, and molecularly and functionally compared them to hPSC-derived melanocytes differentiated through an early NC intermediate^21^. Further, we demonstrated that the transcriptional signature of SCP-derived melanocytes is linked to differences in survival outcomes in melanoma. Finally, by performing a CRISPR screen using an *in vivo* metastasis model, we identified SCP-derived melanocyte transcripts that promote metastasis in melanoma.

## Results

### Emergence of heterogeneity in fate potential during neural crest lineage progression

To systematically characterize how hPSC-NCC populations change over time, we performed single-cell RNA sequencing (scRNAseq) at five stages (S1-S5) of our previously established Schwann cell differentiation^16^. The timepoints sequenced were chosen to capture different stages of NCC specification and maturation: the earliest emergence of SOX10+ NCCs during the induction phase, an early, intermediate, and late stage of neural crest progression, and an early stage of glial induction **(Figure 1A-B)**. Each timepoint was clustered and analyzed separately to preserve real-time information **(Figure 1C)**, and clusters were annotated by the expression of established lineage markers **(Figure S1A)**.

**Figure 1:**
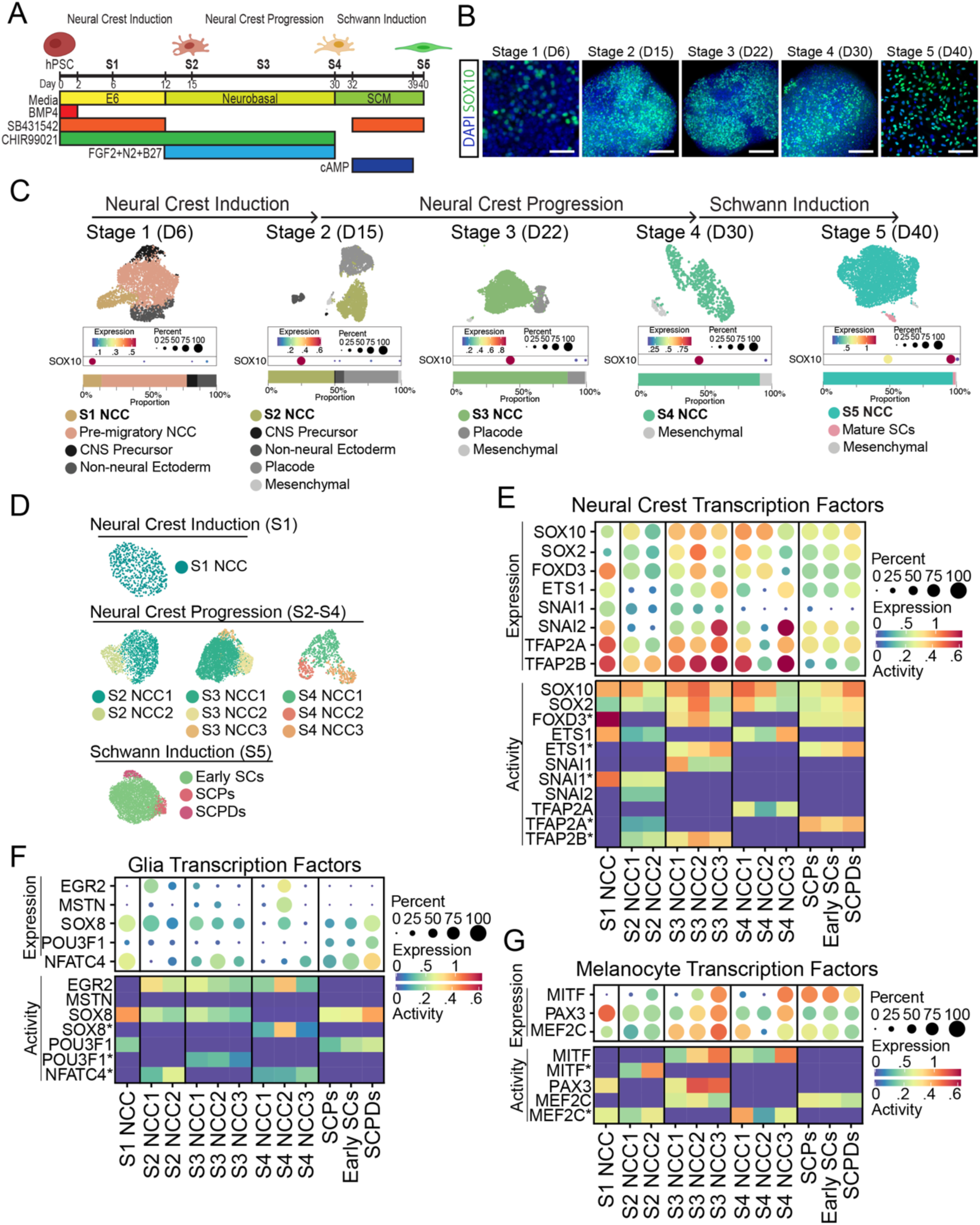
Melanogenic NCCs emerge during hPSC differentiation towards Schwann cells. A. Schematic of directed differentiation protocol to differentiate human pluripotent stem cells to Schwann cell precursors with defined media conditions. The protocol consists of three phases: neural crest induction, neural crest progression and finally Schwann induction. Abbreviations: E6 = Essential 6, SCM = Schwann Cell Media. B. Representative SOX10 expression (green) in hPSC-derived NCCs in stage 1 monolayer neural crest induction, stage 2-4 crestospheres and stage 5 SCP induction. Scale bar represents 100um. C. UMAP embedding, SOX10 expression, cluster proportions, and cluster annotation of stages 1-5 of hPSC neural crest induction, progression, and Schwann induction. D. UMAP embeddings and subclustering of stages 1-5 SOX10+ clusters. E. Expression (top) and predicted activity (bottom) of canonical neural crest transcription factors in stage 1-5 NCC and Schwann subclusters. Asterisks represent TF activity predicted from lower confidence motif annotations. F. Expression (top) and predicted activity (bottom) of canonical glial transcription factors in stage 1-5 NCC and Schwann subclusters. Asterisks represent TF activity predicted from lower confidence motif annotations. G. Expression (top) and predicted activity (bottom) of canonical melanocyte transcription factors in stage 1-5 NCC and Schwann subclusters. Asterisks represent TF activity predicted from lower confidence motif annotations.

On day 6 (D6) of the NC induction, pseudotime analysis revealed that the cell-type composition of the cultures closely resembles the developing neural plate, containing PAX6+/EMX2+ central nervous system (CNS) precursors, GATA3+/CDH1+ non-neural ectoderm (NNE), PAX3+/WNT1+ pre-migratory NCCs, and SOX10+/SNAI2+ stage 1 NCCs emerging from the pre-migratory NCC clusters **(Figure S1B)**. Studies in animal models have identified a complex network of WNT, FGF, and BMP signaling required to pattern the neural plate and for NCCs to develop along the neural plate border^22,23^. Interestingly, the prediction of ligand-receptor interactions using CellChat analysis^24^ suggests these signaling networks establish themselves in human cells *in vitro* at stage 1 of differentiation **(Figure S1C)**. Importantly, the CNS precursor population also scored highly for the expression of negative WNT and BMP gene modules **(Figure S1D)**. Taken together, these data suggest that our directed NC differentiation strategy generates cultures reminiscent of the developing neural plate border to facilitate the signaling necessary for efficient specification/generation of NCCs from hPSCs. This NCC induction strategy resulted in heterogeneous cultures at D15 containing CNS precursors, non-neural ectoderm, SIX1+/POU4F1+ placode, and TWIST1+ mesenchymal cells. These “off-target” but developmentally relevant cell-types were then selected against during the 3D neural crest progression phase, where NCCs selectively aggregated with other NCCs until D30, yielding cultures of >75% SOX10+ cells by day 22 **(Figure 1C and S1E)**.

To characterize the transcriptional changes in our NCCs as they progress through differentiation, we next aimed to analyze the heterogeneity of the SOX10+ NCC clusters across the time points **(Figure 1C)**. The emerging S1 NCCs were homogenous, remaining as a single cluster, while S2, S3, and S4 NCCs contained two, three, and three subclusters, respectively **(Figure 1D)**. Stage 5 NCCs also contained three subclusters **(Figure 1D)**, which were annotated by comparison to our previously published^16^ Schwann cell dataset from a similar differentiation stage. For this comparison we used similarity weighted non-negative embedding (SWNE)^25^. We generated the dimensionally reduced SWNE space based on the published dataset and the stage 5 NCCs were subsequently projected into the same SWNE dimensions, such that transcriptionally similar cells were plotted in the same coordinates (**Figure S1F)**. All subclusters expressed canonical NC transcription factors (TFs) SOX10, SOX2, FOXD3, TFAP2A/B, and SNAI1 or SNAI2, with expression levels varying between clusters but not correlating directly to SOX10 expression **(Figure 1E, top)**.

A new model of NCC fate determination describes the co-activation of competing lineage modules which resolve into fate-biased and then fate-committed NCCs^26^. To determine if hPSC-derived NCC populations exhibit differences in fate potential, we examined both the expression of transcription factors involved in early neuronal (sensory and autonomic), melanogenic, gliogenic, and mesenchymal fate specification, as well as their predicted activity using single-cell regulatory network inference and clustering (SCENIC) analysis^27^. While SOX10 is predicted to be active in all SOX10+ subclusters, with activity level correlating to expression level, other TFs like SNAI2 in S3 and S4 NCCs, TFAP2A in S3 NCCs, and TFAP2B in S4 NCCs showed no predicted activity despite high expression levels **(Figure 1E, bottom)**. The results highlight the importance of considering activity as well as expression for predicting fate potential.

Notably, mesenchymal lineage markers showed minimal expression and activity throughout NCC progression, apart from SOX9 in emerging S1 NCCs **(Figure S1G)**. As well, NCC populations showed no bias toward neuronal fates based on neuronal transcription factor expression or activity during any stage of differentiation **(Figure S1H)**. These results suggest that under our Schwann cell differentiation conditions, NCCs do not activate these developmental programs in the absence of exogenous signals for induction of mesenchymal and neuronal fates.

In agreement with our previous findings^16^, the Schwann cell master regulator EGR2 showed the highest level of expression and activity in S4 NCC2 compared to earlier S1-S3 populations and other S4 NCC subtypes **(Figure 1F)**. Interestingly, S4 NCC2 also expressed MSTN and SOX8 with high predicted SOX8 activity, both markers of NCCs transitioning to a SCP or “hub” transcriptional state recently described by Adameyko and colleagues^26,28^ **(Figure 1F)**. “Hub” cells are multipotent late-stage progenitors in the NC lineage that are primed toward Schwann cells and other late-emerging fates. To further characterize the emergence of the “hub” state in our differentiations, we next examined the expression of the “hub” markers SOX8, ITGA4 and SERPINE2 during NC rogression and Schwann induction **(Figure S1I)**. In addition to SOX8 expression, NCC2 showed the highest expression of ITGA4 and SERPINE2 among S4 NCCs, while expression of all three markers was homogeneous among S5 NCC subtypes **(Figure S1I)**. Interestingly, while S2 NCCs showed moderate to low expression of these markers, we observed a high co-expression of all three markers in a subset of S3 NCCs **(Figure S1I)**. Importantly, scoring the dataset published by Adameyko and colleagues^28^ for the top 100 markers of S4 NCC2 showed cells in the “hub” and the Schwann cell branch to be the most transcriptionally similar populations **(Figure S1J)**. At stage 5, all NCC subtypes, SCPs, Early SCs and SCP derivatives (SCPDs), maintain SOX10, FOXD3, and TFAP2A, as well as gain expression and activity of the key Schwann cell master regulator POU3F1 (OCT6) **(Figure 1E-F)**. Taken together, this data suggests that *in vitro* derived NCCs transition through a “hub”-like transcriptional state over the course of Schwann cell differentiation and recapitulate the temporal progression of the NC lineage *in vivo*.

Intriguingly, we found NCC subclusters with expression and activity of the melanocyte master regulator MITF throughout NCC progression **(Figure 1G)**. We identified a single melanogenic cluster at S2 (S2 NCC2), two melanogenic clusters at S3 (S2 NCC2 and 3), and one melanogenic cluster at S4 (S4 NCC3) **(Figure 1G)**. We validated the emergence of SOX10+/MITF+ cells during NCC progression using immunofluorescence staining **(Figure S1K-L)**. Notably, S5 NCC subclusters also showed high MITF expression, however, MITF activity is inhibited likely due to their exposure to the exogenous NRG1 in the Schwann cell culture media which is known to block melanocyte differentiation **(Figure 1G and S1K-L)**^29^. The largest subclusters of NCCs during NCC progression, NCC1 of S2, S3 and S4 appear to be uncommitted, showing minimal bias toward specific NC derivatives based on expression or activity of lineage-specific transcription factors **(Figure 1E-G and S1G-H)**. Taken together, these results indicate that subsets of hPSC-derived NCCs progress toward glial bias/competency but show melanogenic potential throughout NCC progression. This observation falls in line with lineage tracing experiments demonstrating that late migrating NCCs/SCPs also contribute to melanocytes in the skin during development^19^. However, it remains unclear if melanocytes that emerge during NCC progression are transcriptionally or functionally distinct.

### Developing NCCs give rise to transcriptionally distinct melanogenic progenitors

We hypothesized that the presence of two S3 NCC subclusters with MITF activity could either represent different levels of melanocyte maturation of the same melanocytic lineage, or the emergence and specification of a distinct melanocytic lineage. We therefore aimed to determine the lineage relationships between NCC subclusters and identify features involved in melanogenic differentiation **(Figure 2A)**. We first used Monocle pseudotime analysis^30–32^ to predict lineage progression between NCCs at each stage **(Figure 2B, Figure S2A)**. At D15, uncommitted S2 NCC1 was predicted to transition to melanogenic S2 NCC2. Interestingly at D22, pseudotime predicted a branch point within the uncommitted S3 NCC1 with the melanogenic S3 NCC2 and NCC3 populations at separate ending nodes suggesting divergent specification of these two clusters. Finally, at D30, uncommitted S4 NCC1 is predicted to transition to melanogenic S4 NCC3, as well as gliogenic S4 NCC2, suggesting S4 NCCs retain multipotency towards expected lineages.

**Figure 2:**
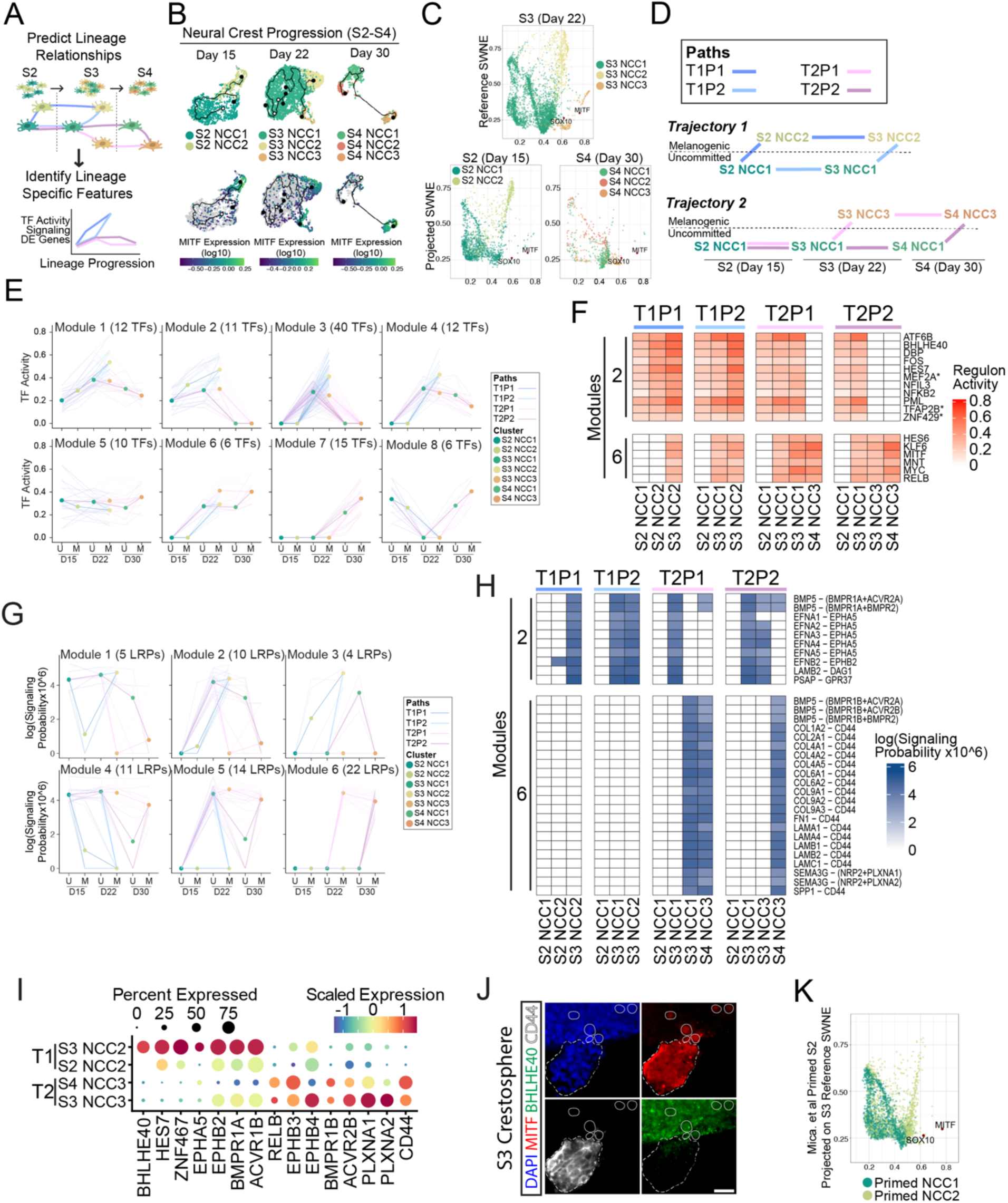
Melanogenic NCCs are predicted to emerge in two distinct trajectories. A. Schematic illustration of melanogenic lineage predictions within hPSC-derived NCC populations and their characterization. B. Top: Pseudotime lineage reconstruction of stage 2-4 neural crest subtypes. Cells (points) are colored by subtype cluster. Black lines follow predicted paths of transcriptional change from selected starting nodes (white circle) to terminal nodes (black circle). Bottom: Log10 MITF expression of stage 2-4 neural crest subtype cells. Grey points represent cells with no detected MITF expression. C. Projection of stage 2 and stage 4 neural crest subclusters onto the SWNE embeddings of stage 3 neural crest subclusters. Cells (points) with similar SWNE coordinates are transcriptionally similar to one another. SOX10 and MITF are projected in the SWNE embedding to show their influence on the cell embeddings. D. Proposed model of lineage “paths” as neural crest subtypes commit to one of two melanocyte differentiation “trajectories.” Trajectory 1 (top) melanogenic NCCs specify from stage 2 or 3 NCCs, while trajectory 2 (bottom) melanogenic NCCs specify from stage 3 or 4 NCCs. Legend abbreviations: T = Trajectory, P = Path. E. Predicted activity of transcription factors along the proposed lineage trajectories and paths. Transcription factors were grouped into 8 modules based on similarity in activity patterns. Modules 1-4 contain transcription factors with higher activity in trajectory 1 melanogenic subtypes, while modules 5-8 contain transcription factors with higher activity in trajectory 2 melanogenic subtypes. Lines are colored by path. Points colored by NC subtype represent the average activity of all transcription factors in the module for that subtype with connected opaque lines to show the average activity pattern. Transparent lines show individual activity patterns for all transcription factors in the module. F. Heatmap of predicted transcription factor activity per stage 2-4 NC subtypes for transcription factor modules 2 and 6. NC subtypes are grouped by proposed lineage path progressions. G. Predicted activity of ligand receptor pairs along the proposed lineage trajectories and paths. Ligand receptor pairs were grouped into 6 modules based on similarity in activity patterns. Modules 1-3 contain ligand receptor pairs with higher activity in trajectory 1 melanogenic subtypes, while modules 4-6 contain ligand receptor pairs with higher activity in trajectory 2 melanogenic subtypes. Lines are colored by path. Points colored by NC subtype represent the average activity of all ligand receptor pairs in the module for that subtype with connected opaque lines to show the average activity pattern. Transparent lines show individual activity patterns for all ligand receptor pairs in the module. H. Heatmap of predicted ligand receptor pair activity per stage 2-4 NC subtypes for ligand receptor pair modules 2 and 6. NC subtypes are grouped by proposed lineage path progressions. I. Scaled expression of trajectory enriched transcription factors and receptors in melanogenic NC subtypes. J. Expression of MITF (red) with T1 progenitor marker BHLHE40 (green) and T2 progenitor marker CD44 (grey) in stage 3 crestospheres. Scale bars represent 25um. K. Projection of stage 2 neural crest subclusters differentiated from the protocol described by Mica et al. onto the SWNE embeddings of stage 3 neural crest subclusters. Cells (points) with similar SWNE coordinates are transcriptionally similar to one another. SOX10 and MITF are projected in the SWNE embedding to show their influence on the cell embeddings.

As there was only a single melanogenic cluster at D30, we next asked which one of the two melanogenic cluster at D22 was maintained through D30. To answer this, we employed two different methods to identify the most transcriptionally similar NCC subclusters between the S2, S3, and S4 NCC subclusters. First, we again utilized SWNE to create a dimensionally reduced space based on the S3 NCC dataset. We then projected S2 and S4 NCC populations into the S3 NCC space, such that transcriptionally similar cells would be plotted in the same coordinates **(Figure 2C)**. Next, we used separate S3 NCC2 and NCC3 transcriptional signatures consisting of the top 100 most significantly differentially expressed genes of each cluster to module score the NCCs of the S2 and S4 samples **(Figure S2B)**. The results from both analyses suggested that the melanogenic S2 NCC2 cluster is closely related to S3 NCC2, while the melanogenic S4 NCC3 cluster was similar to S3 NCC3. Taken together, these data suggest that uncommitted NCCs undergo a temporal transition of differentiating into distinct trajectories of melanogenic progenitors, with S2 NCC2 and S3 NCC2 belonging to trajectory 1 (T1) and S3 NCC3 and S4 NCC3 belonging to trajectory 2 (T2).

Importantly, the pseudotime data showed the potential for the uncommitted NCCs at each given stage to become melanogenic, and the SWNE data suggests that different melanogenic clusters arise from and progress in distinct trajectories. Thus, we propose a model where two paths exist for the differentiation of both T1 and T2 melanogenic progenitors in our culture system. Path 1 of T1 differentiation (T1P1) occurs if uncommitted S2 NCC1 cells differentiate to melanogenic S2 NCC2 which continue to mature directly into melanogenic S3 NCC2 cells by the intermediate time point (D22). Path 2 (T1P2) occurs if uncommitted S2 NCC1 cells maintain potency into the intermediate time point as uncommitted S3 NCC1 cells and then commit to the melanogenic S3 NCC2 identity **(Figure 2D)**. For T2 progenitors, both paths require the uncommitted S2 NCC1 to remain uncommitted as S3 NCC1. Similar to T1, Path 1 of T2 (T2P1) differentiation occurs if uncommitted S3 NCC1 cells differentiate to melanogenic S3 NCC3 cells at D22, which continue to mature directly to melanogenic S4 NCC3 cells by D30. Path 2 of T2 (T2P2) occurs if uncommitted S3 NCC1 cells stay uncommitted until D30 as S4 NCC1 cells and then commit to the melanogenic S4 NCC3 identity **(Figure 2D)**.

Using these predicted differentiation paths, we leveraged SCENIC and CellChat analyses to identify regulons and signaling pathways with differential activities along distinct trajectories. Based on pattern of activity along different paths, transcription factors were grouped into 8 modules **(Figure 2E-F and Figure S2C)**. Module 2 consists of T1 upregulated regulons including HES7 and BHLHE40, while regulons in module 6 such as RELB were upregulated in T2 **(Figure 2F)**. CellChat predicted signaling pathways were grouped into 6 modules **(Figure 2G-H and Figure S2D)**. Interestingly, signaling pathways known to drive melanogenic differentiation such as BMP were shared between the two trajectories, but their activity was predicted to be mediated by different receptors **(Figure 2H and Figure S2D)**. For example, BMP5 was predicted to signal through BMPR1A in T1 (module 2), and BMPR1B in T2 (module 6) **(Figure 2H)**. Other pathways in module 6, like CD44 signaling through the ECM, were specific to T2 progenitors **(Figure 2H)**. Based on these results, we identified a panel of transcription factors and receptors differentially expressed between T1 and T2 melanogenic progenitors **(Figure 2I)**. To validate examples of these T1 and T2 specific markers, we performed immunofluorescent staining and found distinct populations of MITF+ NCCs showing mutually exclusive expression of BHLHE40 and CD44 at D22 **(Figure 2J)**. These analyses revealed the emergence of different populations of progenitors with propensity towards melanocyte differentiation during NCC progression. This prompted us to assess their ability to generate fully differentiated and functional melanocytes.

Studer and colleagues previously established protocols to generate melanocytes from hPSC-derived NCCs utilizing the addition of exogenous BMP4 and EDN3 during the NCC induction phase to prime the early NCCs for melanocyte differentiation^21,33^ **(Figure S2E)**. Given the observation that a distinct melanogenic trajectory emerges from prolonged culture of hPSC-derived NCCs, we sought to compare how “primed” NCCs fit into our two-trajectory model. scRNAseq of the primed culture at S2 (D15) showed an altered cellular composition of off-target cell types, most notably a lack of placode and a larger mesenchymal population, while maintaining a similar percentage of SOX10+ NCCs to our S2 sample **(Figure S2F)**. Similar to the S2 NCCs (D15), sub-clustering of the primed S2 NCCs revealed two subclusters but with a larger proportion of MITF+ cluster NCC2 **(Figure S2G-H)**. Interestingly, when we projected primed S2 NCCs in S3 (D22) NCC SWNE space, we observed an overlap with the uncommitted and T1 melanogenic S3 NCC2 clusters, but not with the T2 progenitor cluster S2 NCC3 **(Figure 2K)**. These analyses suggests that priming increases the induction efficiency of melanogenic NCCs but only produces T1 progenitors. Using the differentiation and maturation conditions described by Studer and colleagues^21,33^, we set out to derive melanocytes from T1 and T2 melanogenic NCC progenitors and compare their molecular and functional properties.

### Early and late NCCs give rise to transcriptionally and functionally distinct melanocytes

Our two-trajectory progenitor model is compatible with the observations *in vivo* that melanocytes differentiate both from early delaminating NCCs as well as from later delaminating NCCs via the nerve-associated SCP intermediate^19,20^. However, early NCC-derived melanocytes and SCP-derived melanocytes are yet to be molecularly and functionally characterized. In order to compare the two populations in a human *in vitro* system, we first had to establish a new differentiation system for generating melanocytes from SCPs.

To achieve this, we subjected both our S5 cultures and primed S2 cultures to the melanocyte induction media developed by Studer and colleagues **(Figure 3A)**. Melanocyte induction of S5 cultures resulted in pigmented cells, here termed T2 melanocytes, which express the canonical melanocyte markers SOX10, MITF, TYR, and PMEL, similar to the melanocytes derived from the primed S2 cultures, here termed T1 melanocytes **(Figure 3B)**. Importantly, both protocols were highly efficient, yielding greater than 90% MITF+ cells **(Figure S3A)**. The primary function of melanocytes is to synthesize melanin within specialized organelles called melanosomes and transfer them to surrounding keratinocytes. We confirmed the ability of both T1 and T2 melanocytes to produce fully mature melanosomes using transmission electron microscopy (TEM) **(Figure 3C)**. To assess melanosome transfer, we set up co-cultures primary human keratinocytes with either T1 or T2 melanocytes. We performed immunofluorescence and flow cytometry for PMEL, a melanosome transmembrane protein, and confirmed the presence of melanosomes in keratinocytes, indicating their successful transfer from both T1 and T2 melanocytes. However, we did not observe any significant differences in transfer kinetics between the two melanocyte populations **(Figure 3D and S3B)**.

**Figure 3:**
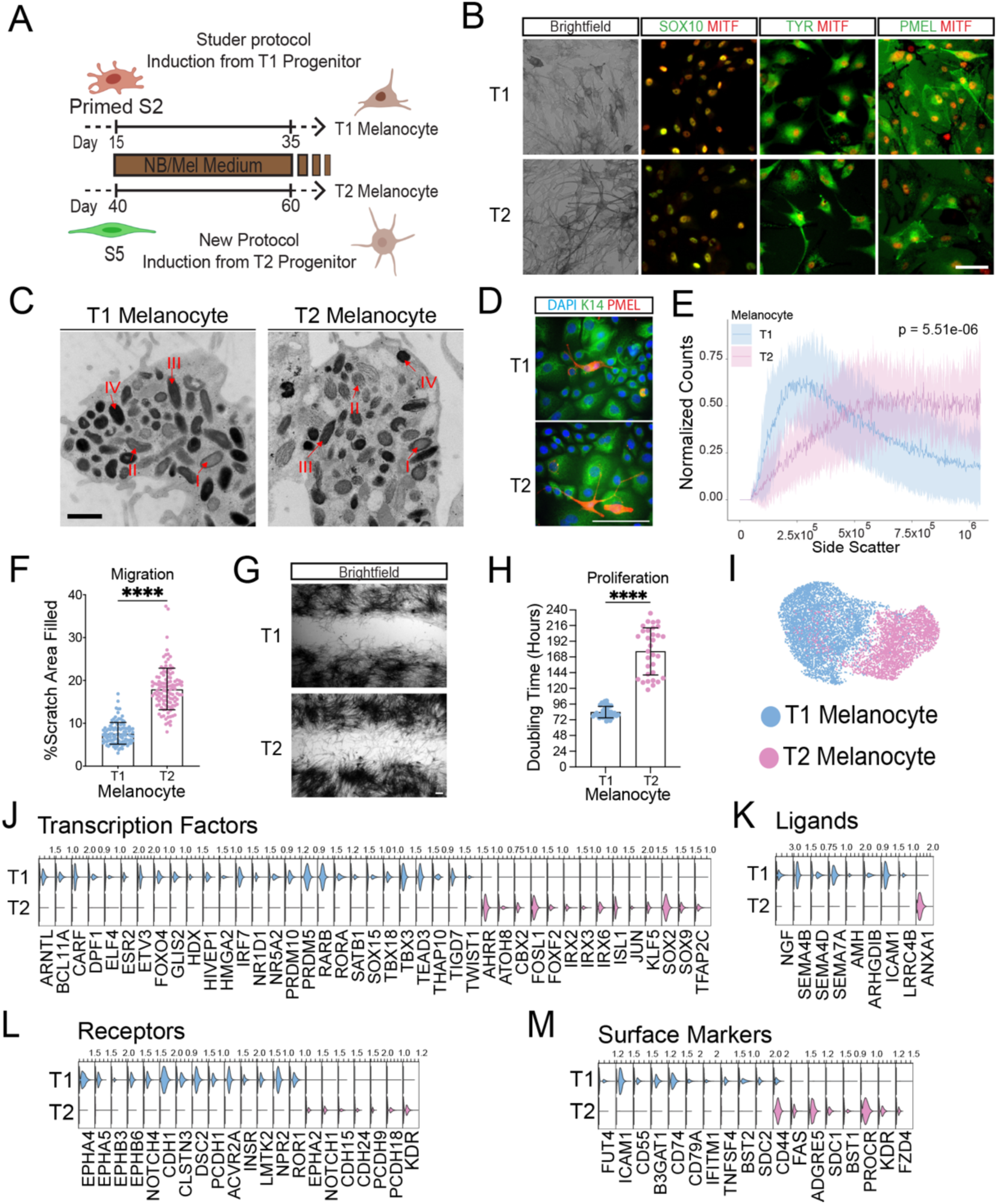
Temporally distinct melanogenic NCCs give rise to functionally and molecularly different melanocytes. A. Schematic of the directed differentiation of trajectory 1 and trajectory 2 melanocytes. Both T1 and T2 melanocytes are differentiated using the same defined medium but differ in that T1 melanocyte differentiation is induced from primed S2 NCCs while T2 melanocyte differentiation is induced from primed S5 SCPs. B. Brightfield images (left) showing pigmentation and immunofluorescent images showing coexpression of MITF (red) with SOX10, TYR and PMEL (green, left to right) in T1 (top) and T2 (bottom) melanocyte cultures. Scale bar represents 100um. C. Transmission electron micrographs of T1 (left) and T2 (right) melanocytes showing melanosome formation. Examples of melanosomes in different parts of the four stages of melanosome maturation are highlighted with red arrows. Scale bar represents 1um. D. Co-culture of T1 (left) and T2 (right) melanocytes with primary human keratinocytes showing transfer of PMEL+ melanosomes (red) to K14+ keratinocytes (green). Scale bar represents 100um. E. Flow cytometry-based quantification of pigmentation via side scatter value. Solid lines show the average distribution of side scatter values with transparent ribbons showing standard deviation (n=12). F. Quantification of T1 and T2 melanocyte scratch assays. Y axis represents the percentage of the original scratch area now covered by melanocyte cell bodies. Points represent technical replicates from n=3. **** is p<0.0001 and error bars represent standard deviation. G. Representative bright field images of T1 (top) and T2 (bottom) melanocyte scratch assays 72hr after scratch. Scale bar represents 100um. H. Doubling time analysis of T1 and T2 melanocytes. Points represent technical replicates from n=3. **** is p<0.0001 and error bars represent standard deviation. I. UMAP embedding of merged T1 and T2 melanocyte datasets. J. Violin plots showing the expression of selected T1 and T2 melanocyte specific transcription factors. K. Violin plots showing the expression of selected T1 and T2 melanocyte specific secreted ligands. L. Violin plots showing the expression of selected T1 and T2 melanocyte specific receptors. M. Violin plots showing the expression of selected T1 and T2 melanocyte specific surface markers.

Interestingly, the quantification of melanin content using flow cytometry and 475nm wavelength absorbance revealed that T2 melanocytes possessed higher pigmentation and melanosome numbers **(Figure 3E and Figure S3C)**. Scratch assays of T1 and T2 melanocytes revealed that T2 melanocytes are also more migratory than T1 melanocytes **(Figure 3F-G)**. However, T1 melanocytes were more proliferative exhibiting a shorter doubling time than T2 melanocytes **(Figure 3H)**. These data suggest that while T1 and T2 melanocytes similarly expressed the canonical melanocytic and pigmentation markers, they showed differences in their functional features.

Given the functional differences between T1 and T2 melanocytes, we next sought to characterize the underlying transcriptional differences by performing scRNAseq on T1 and T2 melanocytes. The two populations were processed and sequenced in a single batch and were age matched based on the time they were cultured in melanocyte induction media. Analysis of both melanocytes individually revealed very homogenous populations, however, integration of these samples results in minimal overlap indicating they are transcriptionally distinct **(Figure S3D and Figure 3I)**. Differential gene expression testing indeed revealed 1557 differentially expressed genes, with 658 genes enriched in T2 melanocytes versus 889 genes enriched in T1 melanocytes **(Figure S3E)**. Of these DE genes, we identified many transcription factors specific to each population, such as the cell cycle regulators FOXO4 and ETV3 in T1 melanocytes and SOX2, SOX9 and TFAP2C in T2 melanocytes **(Figure 3J)**. Interestingly T1 melanocytes exclusively expressed neurogenic factors such as NGF, SEMA4B/D and ICAM1 **(Figure 3K)**. Similar to their distinct progenitor stages, T1 and T2 melanocytes expressed unique receptors from the same receptor families, such as EPHA4/5, EPHB3/6, NOTCH4 and CDH1 in T1 melanocytes and EPHA2, NOTCH1 and CDH15/24 in T2 melanocytes **(Figure 3L)**. Importantly, many surface markers are also enriched in each melanocyte population, such as CD44 in T2 melanocytes, offering potential tools for their prospective isolation using fluorescence activated cell sorting (FACS) **(Figure 3M)**.

### Early and late signatures are expressed in different populations of melanocytes *in vivo*

We next sought to leverage published single cell transcriptomics datasets and embryonic mouse tissue to identify the T1 and T2 molecular signatures in primary melanocytes **(Figure 4A)**. We first integrated the annotated melanocyte clusters from eight datasets collected from multiple developmental stages and anatomical locations^34–40^ **(Figure S4A)**. Scoring the primary melanocyte dataset with separate T1 and T2 melanocyte transcriptional signatures consisting of their top 100 significantly differentially expressed genes showed that the majority of primary melanocytes score highly for one signature or the other **(Figure 4B)**. We annotated the primary melanocytes based on having a higher T1 or T2 melanocyte scoreed and quantified the proportions of T1-like versus T2-like melanocytes in different tissue samples. We found that T1 and T2-like melanocytes were present in all developmental stages **(Figure 4C)**. Interestingly, grouping samples by their anatomical location along the anterior-posterior axis suggested an enrichment of T2-like melanocytes in the anterior and T1-like melanocytes in the posterior regions **(Figure 4C)**.

**Figure 4:**
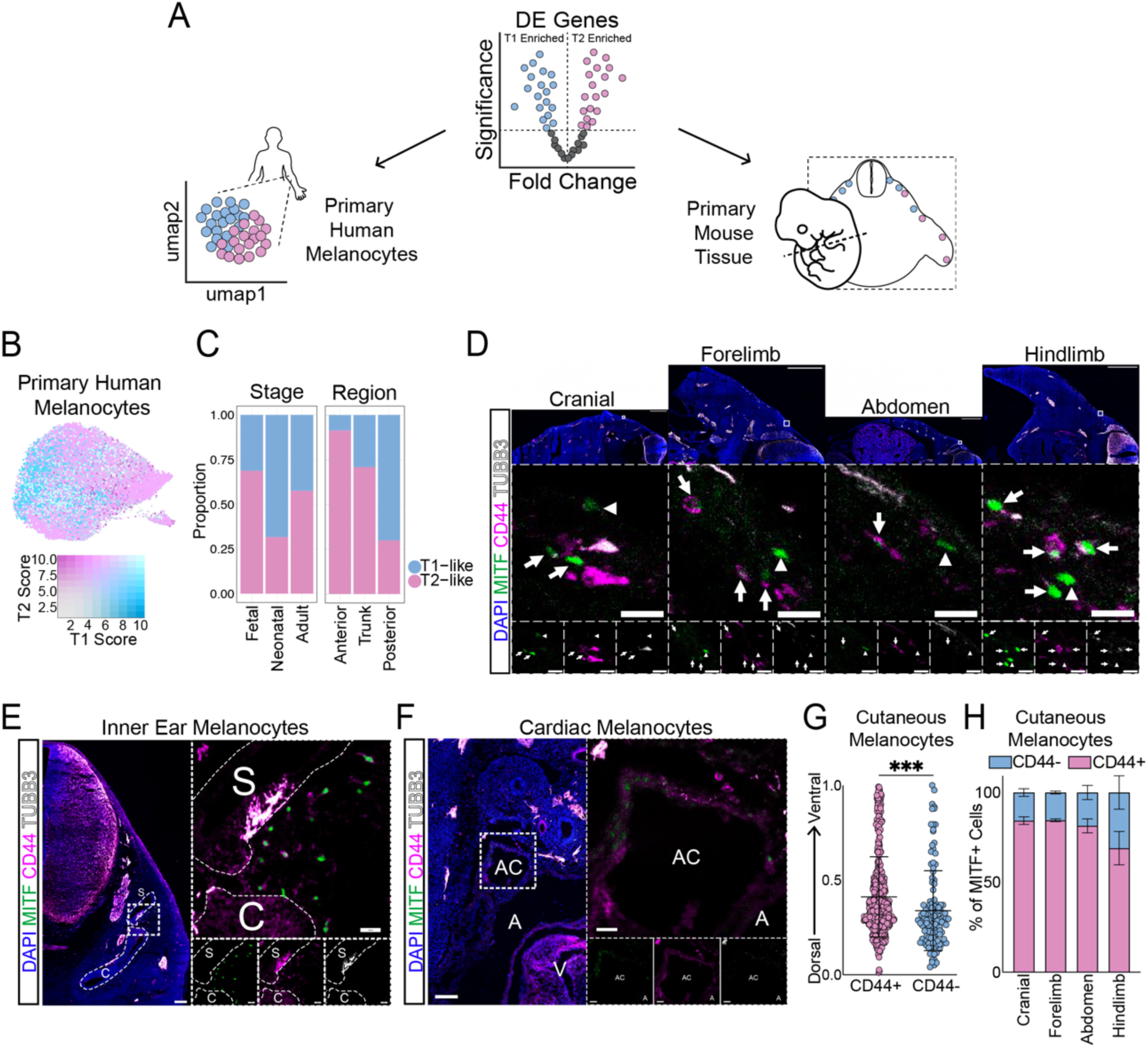
Markers of temporally distinct hPSC-derived melanocytes are expressed by different populations of primary mouse and human melanocytes. A. Schematic illustration of validation of T1 and T2 signature marker in human and mouse primary melanocytes. B. Feature plot of integrated primary melanocytes colored by blended module score for T1 (blue) and T2 (pink) hPSC-derived melanocyte respective top 100 marker genes. C. Bar plots showing the proportion of T1-like and T2-like primary melanocytes grouped by the developmental stage (left) and anterior to posterior anatomical position (right) of the primary melanocytes. D. E12.5 mouse cranial, front limb, abdominal and hind limb sections showing nuclei (blue), melanocytes (MITF, green) neurons (TUBB3, grey) and the T2 melanocyte surface marker CD44 (magenta). Arrows point to CD44+ melanocytes and arrowheads point to CD44-melanocytes. Tile scan scale bars represent 500um, zoomed inlay scale bars represent 25um. E. E12.5 mouse cranial section showing nuclei (blue), melanocytes (MITF, green) neurons (TUBB3, grey) and the T2 melanocyte surface marker CD44 (magenta) around developing inner ear structures (c = cochlea, s = saccule). Tile scan scale bar represents 100um, zoomed inlay scale bars represent 25um. F. E12.5 front limb section showing nuclei (blue), melanocytes (MITF, green) neurons (TUBB3, grey) and the T2 melanocyte surface marker CD44 (magenta) in the developing heart (ac = anterior cardinal vein, a = atrium). Tile scan scale bar represents 100um, zoomed inlay scale bars represent 25um. G. Normalized dorsoventral axis location of CD44- and CD44+ cutaneous melanocytes (n=3 embryos). *** is p<0.001 and error bars represent standard deviation. H. Bar plots showing the proportion of CD44- and CD44+ cutaneous melanocytes grouped by the anterior to posterior anatomical location in sections (n=3 embryos). Error bars represent standard deviation.

Co-staining of MITF with the T2 melanocyte lineage marker CD44 in embryonic day 12.5 mouse cranial, front limb, abdominal and hind limb tissue sections, revealed intermixed populations of CD44+ (T2-like) and CD44-(T1-like) MITF+ cells in all tissue regions **(Figure 4D and S4B)**. In line with observations from SCP-derived melanocyte lineage tracing, we also observed CD44+ MITF+ cells in extracutaneous regions such as the inner ear and heart **(Figure 4E-F)**^41^. Similar lineage tracing of early NC-derived melanocytes and SCP-derived melanocytes also described a dorsoventral bias, such that SCP-derived melanocytes colonize more ventral tissues^19^. Quantification of the relative dorsoventral position of all cutaneous melanocytes across the cranial to hind limb sections similarly showed a significant bias of CD44+ melanocytes to be located more ventral or further from the dorsal neural tube **(Figure 4G)**. To validate the observation of the anteroposterior bias observed in the human primary scRNAseq datasets **(Figure 4C)**, we similarly quantified the number of cutaneous CD44+/MITF+ and CD44-/MITF+ melanocytes across each region and observed a similar trend of a higher proportion of CD44-(T1-like) melanocytes in more posterior tissue **(Figure 4H)**. Taken together, these data provide strong evidence that alternative differentiation trajectories yield distinct melanocyte populations that recapitulate the heterogeneity of melanocytes *in vivo*. Our finding identifies novel markers that enable the isolation and further characterization of these developmentally distinct populations from both mouse and human tissues.

### Signature transcripts of SCP-derived melanocytes promotes metastasis in melanoma

As our data confirmed that T1 and T2 markers were expressed in different subsets of adult melanocytes, we next aimed to understand the implication of the T1 or T2 transcriptional states in melanoma pathogenesis. Based on the functional differences between the two types of melanocytes, we hypothesized that melanomas with a T2-like state may be more migratory and exhibit higher rates of metastasis compared to T1-like melanomas. To test this, we gathered the bulk RNAseq datasets from melanoma samples available on the cancer genome atlas (TCGA) and used the T1 and T2 melanocyte transcriptional signatures to perform gene set enrichment analysis (GSEA) and identify melanoma samples with significantly enriched or reduced expression of these signatures **(Figure 5A, Figure S5A)**. We found that the survival probability of patients with a melanoma enriched for the T2 melanocyte signature was significantly lower than patients with a melanoma reduced for this signature **(Figure 5B, top)**. Conversely, patients with a melanoma enriched for the T1 signature showed higher survival probability compared to those reduced for the T1 signature, although this difference was not statistically significant **(Figure S5B)**. Importantly, direct comparison of patients with a T1-like versus T2-like melanoma showed significantly worse outcomes for T2-like melanoma patients **(Figure 5B, bottom)**.

**Figure 5:**
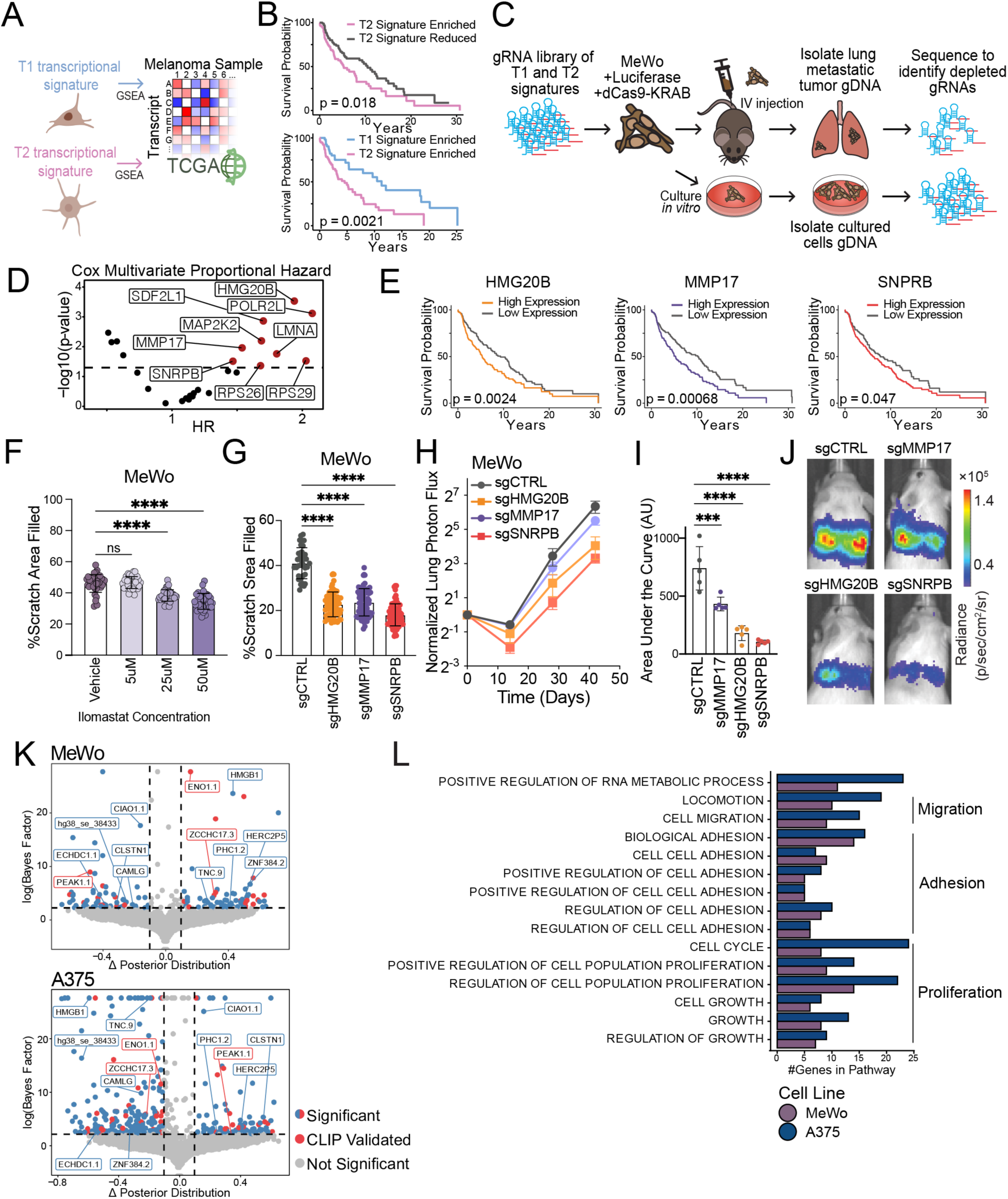
Transcripts enriched in SCP-derived melanocytes promote migration and metastasis in melanoma. A. Schematic illustration of the analysis of 427 melanoma Bulk RNAseq samples from the cancer genome atlas to identify melanomas enriched for T1 and T2 melanocyte transcriptional signatures. B. Top: Survival curves of melanoma patients with melanomas significantly enriched (pink) versus significantly reduced (grey) for the expression of the top 100 T2 melanocyte markers. Bottom: Survival curves of melanoma patients with melanomas significantly enriched for the expression of the top 100 T1 melanocyte markers (blue) versus significantly enriched for the expression of the top 100 T2 melanocyte markers (pink). C. Schematic illustration of the CRISPRi screen for identification of T1 and T2 melanocyte marker genes involved in melanoma metastasis. D. Volcano plot of multivariate Cox proportional hazard ratios versus p-values for CRISPRi hit genes from TCGA melanoma samples. E. Survival curves of melanoma patients with melanomas that highly (colored) versus lowly (gray) express the nominated CRIPSRi hit gene. F. Scratch assay quantification of MeWo cells treated with increasing concentrations of the MMP inhibitor ilamostat. Points represent technical replicates from n=3. p-values are: **** p<0.0001, ns not significant. Error bars represent standard deviation. G. Scratch assay quantification of CRIPSRi MeWo cells transduced with gRNAs targeting the nominated genes versus a non-targeting control. Points represent technical replicates from n=3 assay replicated with n=2 gRNAs. **** is p<0.0001 and error bars represent standard deviation. H. Quantification of luciferase reporter radiance time course in lungs of mice tail vein-injected with CRIPSRi MeWo cells transduced with gRNAs targeting the nominated genes versus a non-targeting control (n=5). Error bars represent standard deviation. I. Area under the curve of luciferase-based lung colonization quantification of *in vivo* tail vein-injected MeWo cells transduced with gRNAs targeting the nominated genes and a non-targeting control (n=5). p-values are: *** p<0.001, **** p<0.0001 and error bars represent standard deviation. J. Representative endpoint images of luciferase reporter radiance in mice tail vein-injected with CRIPSRi MeWo cells transduced with gRNAs targeting the nominated genes versus a non-targeting control. K. Volcano plot of differential splicing analysis between MeWo SNRPB KD cells and Control MeWo cells (top) and between A375 SNRPB KD cells and Control A375 cells (bottom). Solid points are significantly different splicing events (bayes factor >10, difference > .1 or <-.1). Red points are previously validated SNRPB bound transcripts in HEK293T cells. L. Gene Ontology Enrichment analysis of significantly differentially spliced genes in MeWo and A375 SNRPB KD and controls shows enrichment of pathways related to migration and proliferation.

Mortality in melanoma is closely linked to the degree of metastasis. To determine whether T2 melanocyte markers lead to higher metastasis and poor survival, we performed an *in vivo* loss of function screen. For this purpose, we used the MeWo melanoma cell line, which is also significantly enriched for a T2 melanocyte transcriptional signature (Normalized enrichment score = 1.25, adjusted p-value = 0.0018). We used a previously established CRISPR interference (CRISPRi) system^42^ for targeted gene knockdowns, employing a pool of guide RNAs (gRNAs) targeting the 100 genes that make up the T2 signature, as well as the 100 T1 genes as a control set. We transduced MeWo cells that constitutively express dCas9-KRAB and luciferase with the gRNA-expressing lentiviruses in a pooled fashion and injected them into the tail veins of NSG (NOD scid gamma) mice for lung colonization assays. At the experimental endpoint, we isolated the melanoma tumors formed in lungs and sequenced their genomic DNA to determine their gRNA pool. We compared the gRNA abundance in the tumors with transduced MeWo cells grown in parallel *in vitro*, reasoning that any gRNA significantly depleted in the melanoma tumors *in vivo* is likely to target a gene essential for the metastatic process **(Figure 5C)**. In total, gRNAs targeting 39 genes were significantly depleted in lung tumors, of which 66% targeted T2 signature genes **(Figure S5C-D)**.

To prioritize genes for further characterization, we utilized the TCGA melanoma datasets to filter hits in two steps. We performed a multivariate Cox proportional hazard test which first determined whether the hit gene’s expression level predicted a significant difference in survival probability. Then, we considered additional variables such as patient age, treatments received and stage at diagnosis to determine if the difference in survival probability based on the gene’s expression levels remained significant. Nine genes passed the criteria, and of these we selected HMG20B, MMP17, and SNRPB for further validation experiments **(Figure 5D-E)**. While HMG20B and MMP17 have been shown to be highly expressed in melanomas^43^, they have never directly been implicated in melanoma metastasis. However, both HMG20B, MMP17 and other members of these gene families have been implicated in metastasis in other cancers^44–51^. Similarly, SNRPB, a canonical subunit of the spliceosome, has not previously been identified as a melanoma risk factor, but has been implicated in the metastatic process of other cancers like glioblastoma, cervical and non-small cell lung cancer^52–54^.

To validate the role of these genes in melanoma metastasis we used a combination of genetic and pharmacological approaches in both the MeWo and A375 cell lines. A375 is another melanoma cell line enriched for the T2 signature (Normalized enrichment score = 1.33, adjusted p-value = 1.99e-05). MMPs are a druggable protein family, and MMP17 is a predicted target of the broad spectrum MMP inhibitor ilamostat. Scratch assays of MeWo and A375 cells treated with increasing concentrations of ilamostat showed a dose dependent decrease in cell migration highlighting the promigratory role of MMPs **(Figure 5F and S5E-F)**. We next generated stable knockdown cell lines using CRISPRi, targeting HMG20B, MMP17 and SNRPB in both MeWo and A375 cells to evaluate the effect on migration and proliferation **(Figure S5G)**. Knockdown of all three genes significantly decreased cell migration, while only knockdown of MMP17 in MeWo cells significantly increased the doubling rate by 3 hours **(Figure 5G and S5H-I)**. Importantly, tail vein injections of MeWo cell lines with stable knockdown of all three genes showed a significant reduction in formation of lung tumors **(Figure 5H-J)**. Notably, the effect sizes of the lung colonization assay matched with the effect size of the *in vitro* migration assay.

As the knockdown of the splicing factor SNRPB resulted in the greatest delay in lung colonization *in vivo* **(Figure 5H-J)**, we next sought to determine the spliceosome network regulated by SNRPB in melanoma. We performed paired end bulk RNAseq on control and SNRPB knockdown MeWo and A375 cells to identify differentially spliced transcripts. Interestingly, only 13 transcripts were identified as shared differential splicing events in both MeWo and A375 SNRPB knockdown cells **(Figure 5K)**. Notably, many unique differential splicing events were validated as SNRPB targets in a CLIP dataset generated in HEK293T cells **(Figure 5K)**. Intriguingly, some of the differentially spliced transcripts in both MeWo and A375, such as HMGB1 and PEAK1, showed opposite trends in transcript isoform abundance **(Figure S5K-L)**. Importantly, despite the minimal overlap in differentially spliced transcripts, gene ontology analysis of all differentially spliced genes in both the MeWo and A375 lines revealed that these genes belong to many shared biological process pathways relevant to cell migration, adhesion and proliferation **(Figure 5L)**. Taken together, these data suggest that both the transcript targets of SNRPB and the isoform prevalence altered by SNRPB were cell line specific. However, in a melanoma background SNRPB collectively influenced metastasis through the alternative splicing of transcripts involved in migration, adhesion and proliferation.

## Discussion

Our study provides a comprehensive temporal map of human NCC lineage specification in a hPSC-derived model. Profiling NCCs in high temporal resolution has been challenging to achieve, particularly in humans, due to ethical and technical limitations associated with obtaining fetal tissue. Newly advanced hPSC differentiation systems recapitulate the process of embryonic and fetal development with high precision and provide access to transient populations such as NCC. For example, single cell profiling at early stages of our differentiation revealed cell populations and predicted signaling interactions that mirror that of the developing neural plate. Using an *in vitro* induction system that mimics the normal developmental environment captures complex aspects of NCC fate specification that replicates *in vivo* conditions. Overall, our results highlight the power of this hPSC-based system in studying aspects of neural crest specification, differentiation and identity. Further characterizations of hPSC-NCCs, including profiling their epigenetic states, will be necessary to fully compare them to analogous mouse and chick populations.

Our transcriptomic profiling of differentiating hPSCs at single cell resolution revealed dynamic and increasing transcriptional heterogeneity within the NCC populations over time. Lineage analysis of NCC subtypes revealed a sustained population of uncommitted NCCs which maintain melanogenic competency. Interestingly, the melanogenic progenitors produced by stage 2 and stage 4 NCCs are transcriptionally distinct, which we refer to T1 and T2 progenitors respectively. In the stage 3 NCCs we captured a transition point where the uncommitted NCCs are competent to produce both melanogenic progenitors prior to becoming restricted to only specifying the T2 progenitors. The kind of melanogenic progenitor produced appears to be tightly linked to the age of the NCCs, as we showed that existing protocols that included exogenous melanogenic signals such as BMP4 and EDN3 prior to stage 2 do not produce another transcriptionally distinct melanogenic progenitor, but are rather more efficient at producing the T1 melanogenic progenitors. Emergence of these temporally distinct melanogenic NCC is consistent with previous *in vivo* studies that describe two waves of melanogenesis from early NCCs that migrate dorsolaterally through the mesenchyme and late NCCs that migrate ventrolaterally along the developing nerves^19,20^.

Our comparison of T1 and T2 progenitors identified 114 transcription factors and 70 ligand-receptor pairs with differential activity patterns. Interestingly, many transcription factors were expressed by both melanogenic progenitors but were predicted to be differentially active. This suggests that additional regulatory mechanisms are responsible for the ultimate gene expression differences between T1 and T2 progenitors. Remarkably, T1 and T2 progenitors showed differential expression of genes relevant to their *in vivo* features. For example, T1 progenitors expressed EPHB2 which is necessary for dorsolateral migration of NCCs and melanoblasts, while T2 progenitors expressed CD44, a neuronal cell adhesion molecule utilized by glia for nerve track migration. This highlights the utility of hPSC models for the study of developmental processes.

We show high efficiency derivation of pigmented melanocytes from T2 progenitor that exhibit higher levels of pigment production. Interestingly, both types of melanocytes are capable of transferring their melanosome to keratinocytes. The physiological implication of this pigmentation difference remains to be determined and possible links between the distribution of T1 and T2 melanocytes in different skin regions and levels of pigment production should be explored further. While T1 and T2 melanocytes expressed the same canonical markers, they were largely transcriptionally unique, with greater than 1500 differentially expressed genes. Importantly, we used these transcriptional signatures to identify T1-like and T2-like melanocytes in published human scRNAseq datasets and mouse fetal tissue, showing that both kinds of melanocytes persisted into adulthood and their markers are conserved. Consistent with their predicted ventrolateral migration path^20^, T2-like melanocytes were more ventrally distributed within the fetal skin. Interestingly, T1-like melanocytes were more abundant in posterior regions both in fetal mouse tissue and in adult human skin samples. Additionally, the transcriptomic comparison points to their potentially unique functions aside from melanin production and transfer. T1 melanocytes uniquely expressed NGF and semaphorins suggesting a role in neurotrophic support and axon guidance. Future studies will be required to validate this developmental role specific to T1 melanocytes and may provide new etiological insights into neurocristopathy syndromes involving pigmentation defects and peripheral neuropathies.

We found that T2 melanocytes migrate faster than T1 melanocytes, and a T2 signature in melanomas is associated with lower survival probability possibly due to a higher metastatic potential. Our *in vivo* CRISPRi screen revealed that of the 100 most significantly enriched genes in T2 melanocytes, 25% played a role in metastatic lung colonization in mice. For 9 of these genes, including HMG20B, MMP17 and SNRPB, expression level alone was a significant predictor of melanoma patient survival outcome. Knockdown of these three transcripts in melanoma cells was sufficient to decrease both migration and lung colonization, with SNRPB knockdown resulting in the most substantial reduction of lung colonization. SNRPB is an RNA splicing factor, and alternative splicing events have previously been identified as markers of oncogenic progression in melanoma^55,56^. Although SNRPB was previously identified as an oncogenic candidate in glioblastoma^52^, its possible roles in melanoma were not identified. Our data suggest that SNRPB controls the splicing of transcripts involved in in migration and adhesion of melanoma cells. Future work will be necessary to mechanistically understand how the alternatively spliced isoforms regulate these cellular functions. Collectively, these results indicate that melanomas that display the unique signatures of melanocytes from temporally distinct developmental origin constitute different categories of melanoma and likely require personalized therapeutic approaches. Future studies will be necessary to show direct oncogenic transformation of each type of melanocyte and the conservation of these original transcriptional signatures during the tumorigenesis process. Nonetheless, the presence of transcriptionally T1-like and T2-like melanocytes into adulthood offer new insight into origins of heterogeneity in melanoma cases.

In conclusion, our findings highlight the utility of directed stem cell differentiations as models of human neural crest development and provide a platform for developing in-depth mechanistic understandings of NC diversification, cell fate decisions, lineage progression and restriction. Furthermore, our new T2 melanocyte differentiation strategy allows for the first comparison of NC-derived and SCP-derived melanocytes and provides more comprehensive *in vitro* melanocyte models necessary for disease modeling and drug discovery. Finally, our work proposes developmental origin as a new source of transcriptional and functional heterogeneity and survival outcomes in melanoma and provides a basis for selection of origin specific gene targets for personalized treatment.

## Acknowledgements

We thank the UC Davis Biological Electron Microscopy Core Facility for their technical support and sample preparations. We acknowledge the SF Chan Zuckerberg Biohub Sequencing platform with special thanks to Norma Neff, as well as the UCSF Center for Advanced Technology for next generation sequencing. We thank the Laboratory Animal Resource Center (LARC) at UCSF. The work was generously supported by grants from UCSF Program for Breakthrough Biomedical Research and Sandler Foundation, the NIH Director’s New Innovator Award (DP2NS116769) and the National Institute of Diabetes and Digestive and Kidney Diseases (R01DK121169) to F.F. The study was supported in part by UCSF Laboratory for Cell Analysis shared resource facility through the NIH grant P30CA082103 and by grants to H.G. from the National Cancer Institute (R01CA24098 and R01CA244634). A.N. was supported by DoD PRCRP Horizon Award W81XWH-19-1-0594. E.G. was supported by 1F31HD108875 and D.L. by 1R01ES028212 and 1R01GM122902.

## Author contributions

R.S. Designed, performed, and analyzed all *in vitro* experiments including flow cytometry, 2D and whole-mount immunofluorescent staining, qPCR, co-cultures, melanosome transfer, melanin quantification, doubling rate, and migration assays. Designed and executed all single cell RNA sequencing and TCGA data analysis. Performed analysis of embryonic mouse images. Performed analysis and generated figures of CRISPRi screen. Performed analysis and generated figures of spliceofrom data.

A.N. Assisted and supervised the generation of all melanoma CRISPRi line, generation of the CRISPR gRNA library, the *in vivo* CRISPRi screen and performed bulk RNA sequencing library preparations.

A.M Performed 10X library preparations for scRNAseq and assisted with scRNAseq data processing and analysis.

E.G. Performed dissection, sectioning, immunofluorescent staining and imaging of mouse embryos.

K.G. Performed injections for mouse experiments, assisted with luciferase imaging of CRISPRi screen, and performed mouse tissue collection and processing.

H.M. Assisted with *in vitro* sample collection.

M.R. Assisted with *in vitro* sample collection.

N.E. Assisted with figure generation and design.

D.L. Assisted with scRNAseq data processing and analysis.

P.N. Assisted with *in vitro* arm of CRIPSRi screen, assisted with CRISPRi line validation.

B.S. Performed transmission electron microscopy sample preparation and supervised transmission electron microscopy imaging.

M.L.L. Performed mouse husbandry, breeding, and embryo collection.

L.S. Supervised mouse husbandry, breeding, and embryo collection.

D.L. Supervised mouse embryonic tissue staining and imaging.

S.D. Supervised 10X scRNAseq sample collection and sequencing.

H.G. Supervised all CRISPRi experiments. Supervised and performed mouse injections. Performed bulk RNAseq spliceoform analysis.

F.F. Designed and conceived the study, supervised all experiments, and wrote the manuscript.

## Tables

Table 1. Antibodies and dilutions used for immunofluorescent staining

Table 2. scRNAseq dataset quality control filtering metrics

Table 3. scRNAseq dataset clustering parameters

Table 4. Cluster transcriptional signature used for module scoring

Table 5. Primers and oligo sequences used in CRISPRi experiments

## Methods

### Maintenance and passaging of human pluripotent stem cells (hPSCs)

The H9 human embryonic stem cell (hESC) line was maintained on Geltrex (Gibco) coated plates in chemically defined media (Essential 8, Gibco) at 37°C 5% CO_2_. Stem cell media was changed every other day and stem cell colonies were passaged at ∼70% confluency using EDTA (Corning) to dissociate. Stem cell maintenance cultures were tested for mycoplasma contamination once a month.

### Maintenance and passaging of melanoma cell lines

MeWo cells were grown in EMEM (Lonza) supplemented with 10% FBS (ScienCell), penicillin and streptomycin (Gibco) and amphotericin B (Gibco). A375 cells were grown in DMEM supplemented with 10% FBS, penicillin, streptomycin and amphotericin B. Media was changed every other day and cultures were passaged at 90% confluency using 0.05% Trypsin (Corning). Cultures were tested for mycoplasma contamination once a month.

### Cranial neural crest induction from hPSCs

hESC cultures at ∼70% confluence were dissociated using EDTA and replated one to one on Geltrex coated plates in stem cell media (Essential 8) to establish an hESC monolayer. The following day, cranial neural crest induction was initiated (D0) by replacement of the stem cell maintenance media with neural crest induction media A (Essential 6 base medium (Gibco), 600nM CHIR99021 (Biogems), 10uM SB431542 (Selleckchem), and 1ng/ml BMP4 (R&D Systems)). On days 2, 4, 6, 8, and 10 of cranial neural crest induction cultures are fed with neural crest induction media B (Essential 6 base medium, 1.5uM CHIR99021, 10uM SB431542). On day 12, cranial neural crest induction cultures are dissociated with Accutase (Innovative Cell Technologies) for 20min, at 37°C with 5% CO_2_, pelleted at 300xg for 2min, resuspended in neural crest maintenance media (NC-C) (Neurobasal base medium (Gibco), 20ul/ml B27 supplement (Gibco), 10ul/ml N2 supplement (Gibco), 10ul/ml Glutagro (Corning), 10ul/ml MEM NEAAs (Corning), 3uM CHIR99021 and 10ng/ml FGF2 (R&D Systems)) and plated in ultra-low-attachment plates (Corning). This culture format selects against contaminating ectodermal lineages that arise during the neural crest induction and maintains neural crest cells in 3D crestospheres. On days 14, 18, 22, and 26 the crestospheres are fed with fresh NC-C media by gently swirling the plate on a flat surface to collect the spheres in the center of the well, aspirating the spent media carefully from the edges of the well without removing the crestospheres and replacing with fresh NC-C. To prevent spontaneous differentiation in the center of the crestopsheres, crestosphere cultures are passaged on days 16, 20, 24 and 28. The crestopsheres are collected and centrifuged at 300xg for 1min and resuspended in Accutase. Incubation time in Accutase starts with 15min on D16 and increases by 5min each dissociation day as neural crest maintenance progresses to achieve full dissociation. Once dissociated, the neural crest cell suspensions are pelleted at 300xg for 2min and resuspended in fresh NC-C and plated in a new ultra-low-attachment plate.

### Schwann cell induction from hPSC-derived neural crest

On day 30 of the cranial neural crest cultures, crestopsheres are dissociated with Accutase as described above for 30min at 37°C with 5% CO_2_. After centrifugation at 300xg for 2min, the cell pellet is resuspended in serum-free Schwann Cell Medium (SCM, ScienCell) and plated on poly-L-ornithine (PO, Sigma)/Fibronectin (Sigma)/Laminin (R&D Systems) coated plates. During Schwann induction, fresh SCM (serum-free) is fed daily. Starting on day 32 until day 38, SCM is supplemented with 100um cyclic AMP (cAMP, Sigma) to promote Schwann cell differentiation and 10uM SB431542 to prevent off-target differentiation. At day 39, cAMP is no longer added to the medium. Cultures were maintained in serum-free SCM and passaged at 90% confluence using 0.05% Trypsin for 1min at 37°C, with 5% CO_2_.

### Melanocyte primed neural crest induction and T1 melanocyte differentiation from hPSCs

#### From Studer and colleagues

hESC cultures at ∼70% confluence were dissociated using EDTA and replated one to one on Geltrex coated plates in stem cell media (Essential 8) to establish an hESC monolayer. The following day, cranial melanocyte-primed neural crest induction was initiated (D0) by replacement of the stem cell maintenance media with neural crest induction media A (Essential 6 base medium, 600nM CHIR99021, 10uM SB431542, and 1ng/ml BMP4). On days 2 and 4 cultures are fed with neural crest induction media B (Essential 6 base medium, 1.5uM CHIR99021, 10uM SB431542). On days 6, 8, and 10, cultures are fed with melanocyte priming media (MPM) (media B with 10ng/ml BMP4 and 100ng/ml EDN3 (Sigma)) to bias the cranial NCCs towards a melanocyte fate. On day 12, primed neural crest induction cultures are dissociated with Accutase for 20min at 37°C with 5% CO_2_, pelleted at 300xg for 2min, resuspended in neural crest maintenance media (NC-C) (Neurobasal base medium, 20ul/ml B27 supplement 10ul/ml N2 supplement, 10ul/ml Glutagro, 10ul/ml MEM NEAAs, 3uM CHIR99021 and 10ng/ml FGF2) and plated in ultra-low-attachment plates. On day 14 the primed crestospheres are fed with fresh NC-C media by gently swirling the plate on a flat surface to collect the spheres in the center of the well, aspirating the spent media carefully from the edges of the well without removing the crestospheres and replacing with fresh NC-C. On day 15 the primed crestopsheres are collected and centrifuged at 300xg for 1min and resuspended in Accutase for 15min at 37°C with 5% CO_2_ to dissociate. Once dissociated, the primed neural crest cell suspensions are pelleted at 300xg for 2min and resuspended in melanocyte induction media (MIM) (50% Neurobasal, 30% Low glucose DMEM/F12 (Gibco), 20% MCDB201 base medium (Sigma), 2% B27 supplement, 1% Glutagro, 0.8% ITS+ (Gibco), 25ng/ml BMP4, 100ng/ml EDN3, 50ng/ml SCF (Peprotech), 4ng/ml FGF2, 50ng/ml Cholera Toxin Beta Protein (Novus), 3uM CHIR99021, 100uM Ascorbic Acid (Sigma), 500uM cAMP, 50nM Dexamethasone (Sigma)) and plated on poly-L-ornithine (PO)/Fibronectin/Laminin coated plates. Cultures are fed with fresh MIM every other day. Cultures are maintained at confluence or passaged for expansion by dissociation with Accutase for 5-10min at 37°C with 5% CO_2_.

### T2 Melanocyte induction from hPSC-derived schwann cell precursors (SCPs)

Following differentiation to Schwann cell precursors to Day 40, cultures were switched to MIM (used above). Cultures are fed with fresh MIM every other day. Cultures are maintained at confluence or passaged for expansion by dissociation with Accutase for 5-10min at 37°C with 5% CO_2_.

### 2D Immunofluorescence

The adherent culture’s media was aspirated, and cells were washed with PBS (Gibco). After washing, cells were fixed with 4% PFA (Santa Cruz) for 30 min at room temperature. After fixation cells were washed 3x with PBS prior to permeabilization using the BD Perm/Wash buffer 1X permeabilization buffer (PB, eBiosciences) for at least 30min at room temperature. Primary antibodies were diluted (Table 1) in PB and incubated on cells overnight at 4°C. Cells were washed with PB 3x for 5min each and incubated with secondary antibodies (Invitrogen) diluted 1:1000 in PB for 1hr at room temperature. Cells were again washed with PB 3x for 5min and finally placed in PBS for imaging on an epi-fluorescent microscope (Echo Revolve).

### Whole mount Immunofluorescence

Crestospheres were transferred to Eppendorf tubes, centrifuged at 200xg for 1min (same parameters used for all subsequent centrifugation steps) and washed with PBS (all incubations and washes done on orbital rocker). Crestospheres were fixed with 4% PFA for 30minutes at room temperature and then washed with PBS 3x for 5min. Crestospheres were then permeabilized with 0.5% Triton X-100 (Thermo) in PBS for 20min at room temperature and then blocked with blocking buffer (0.1% Triton X-100, 5% donkey serum (Jackson Labs), and 1% BSA (Sigma) in PBS). Primary antibodies were diluted (Table 1) in blocking buffer and incubated with crestospheres for 48hrs at 4°C. Crestospheres were washed with blocking buffer 3x for 20min and then incubated with secondary antibodies diluted 1:1000 in blocking buffer for 48hrs at 4°C. Crestospheres were again washed with blocking buffer 3x for 20min and finally placed in PBS in coverslip chamber slides for confocal imaging using the molecular devices imageXpress microscope.

### Flow Cytometry

Single cell suspensions were made using the passaging methods listed above depending on the time point being collected. Cells were pelleted by centrifugation at 500xg for 3min (same parameters used for all subsequent centrifugation steps) and washed with PBS. Cells were then fixed using the FOXP3/Transcription Factor staining buffer set 1X fixation/permeabilization buffer (eBiosciences) for 30min at 4°C. Cells were washed with PBS and then resuspended in 1X PB for 30min at room temperature. Primary antibodies were diluted (Table 1) in PB and incubated on cells overnight at 4°C. Cells were washed with PB 3x for 5min and incubated with secondary antibodies diluted 1:1000 in PB for 1hr at room temperature. Cells were again washed with PB 3x for 5min and finally resuspended in PBS and transferred to FACS tubes for cytometry analysis using an Attune NxT (Thermo). Population percentage-based analysis was performed using Flowjo (v10). See “melanin quantification” below for side-scatter analysis methods.

### Quantitative reverse transcription PCR (RT-qPCR)

RNA was extracted using the PureLink RNA Isolation kit (Thermo). Briefly, cells were dissociated as described above, pelleted and resuspended in lysis buffer. Cell lysis suspensions were vortexed, snap frozen and thawed prior to column purification and elution in water. RNA stock concentrations were quantified by nanodrop. cDNA was generated using qSCRIPT cDNA SuperMix (Quantabio) using 1ug of RNA as input. cDNA was diluted to approximately 10ng/ul and used as input for 10ul qPCR reactions containing 10ng cDNA, 2.5uM primers (Table 5) and 50% PowerUp™ SYBR™ Green Master Mix (Thermo). qPCR reaction amplifications were measured using a QuantStudio 6 Real Time PCR system (Thermo).

### Transmission electron microscopy sample preparation and imaging

Melanocytes were dissociated as described above and counted using a hemocytometer. Melanocytes were plated 10,000 cells/cm^2^ in a PO/Fibronectin/Laminin coated permanox plastic Nunc™ Lab-Tek™ chamber slides (Thermo) in MIM. Melanocytes were cultured in slide chambers for 24hrs before fixation with a modified Karnovsky’s fixative (2% formaldehyde, 2.5% glutaraldehyde, and 0.1M sodium phosphate, EMS) for 24hrs at 4C. Cells were washed with 0.1 M sodium phosphate buffer and then incubated in secondary fix (1% osmium tetroxide and 1.5% potassium ferrocyanide in 0.1M sodium phosphate) for 1 hour. Cells were washed with cold ddH2O 3 times before stepwise dehydration through an ethanol gradient at 30%, 50%, 70%, 95%, and 3×100% ethanol for at least 10 minutes each. Ethanol was removed and 100% resin was added and allowed to infiltrate overnight at room temperature. Next, as much resin as possible was removed from the cells and fresh resin was added (Dodecenyl Succinic Anhydride, Araldite 6005, Epon 812, Dibutyl Phthalate, Benzyldimethylamine). The resin was polymerized overnight at 70C. Resin blocks were sectioned on Leica EM UC6 ultramicrotome at approximately 100nm, collecting sections on copper grids. Grids were dried in an oven at 60C for 20 minutes. Grids were stained with 4% aqueous uranyl acetate and 0.1% lead citrate in 0.1N NaOH. Sections were imaged in a FEI Talos L120C TEM at 80kv.

### Melanocyte-Keratinocyte co-culture and melanosome transfer assay

Primary adult human keratinocytes (ATCC) were cultured on PO/Fibronectin/Laminin coated in Keratinocyte SFM (Gibco). At the start of co-culture, keratinocytes were dissociated with 0.05% Trypsin and counted with Trypan blue using a CellDrop FL (Denovix), while melanocytes were dissociated with accutase as described above and counted using a hemocytometer. Melanocytes and keratinocytes were mixed at a ratio of 1:10 to mimic the estimated ratio observed in human skin^57^, i.e. keratinocytes were seeded at 150,000 cell/cm^2^ and melanocytes at 15,000 cell/cm^2^ in the same well, and plated on PO/Fibronectin/Laminin coated plates. Time 0 was considered to be two hours after initial plating to allow for cell attachment. Co-cultures are collected using 0.05% Trypsin to obtain single cell suspensions processed for Flow cytometry as described above at 24hr, 48hr and 72hrs. Additionally Co-cultures were fixed at 72hrs for 2D immunocytochemistry as described above. Flow and immunocytochemistry samples were stained with K14 (Thermo, Table 1) to mark keratinocytes and PMEL (Thermo, Table 1) as a melanosome incorporated protein. Flowjo was used to analyze melanosome transfer as the percentage of PMEL+/K14+ keratinocytes versus total keratinocytes.

### Bulk quantification of melanin

Melanocytes were dissociated with Accutase as described above and counted using a hemocytometer. Melanocytes were plated at 150,000 cell/cm^2^ in a PO/Fibronectin/Laminin coated plate in MIM. Melanocytes were cultured for 72hrs and then lysed with 1X RIPA buffer (Sigma) using the supplier protocol for lysing of adherent cultures. Briefly, ice cold RIPA buffer was added to the wells of the plate and the plate was incubated on ice for 5min. The RIPA solution was pipetted up and down repeatedly in the well to lyse residual cells and then transferred to a microcentrifuge tube. Cell lysates were centrifuged at 14,000xg for 15 minutes at 4°C. This centrifugation step pelleted all melanin while the DNA remained in the supernatant. The supernatant was then transferred to a new microcentrifuge tube and the pellet containing the melanin is further dissolved by incubation in 250ul of 1M NaOH (Sigma) for 40min at 37C. Once fully dissolved, the lysate was transferred to a 96 well plate and the OD475 was measured using a plate reader (Molecular Devices). In parallel, the RIPA lysate was diluted 1:50 in PBS and mixed with propidium iodide (PI, Thermo) in PBS at a final ratio of 1:3000. PI fluorescence intensity, representing DNA abundance/concentration, was measured by excitation at 535nm and emission detection at 615nm, the maximal excitation/emission of PI when bound to DNA, using a plate reader. All lysates were measured in duplicates and averaged per biological replicate. The OD475 of each pellet dissolved in NaOH was normalized to the PI fluorescence intensity of the matching lysate.

### Single cell quantification of melanin

Melanocytes were prepared for Flow cytometric analysis as described above. Side scatter voltage was set using a 1:1 mixture of T1 and T2 melanocytes. The resulting FCS files were analyzed in R, using the flowCore package (v2.8) for data importation. Each file, representing a biological replicate (BR) of the respective differentiation, was “gated” to remove debris events by filtering for events with a FSC-A value greater than 190000 and a SSC-A value greater than 50000. Each BR’s SSC-A values were then merged into a single event-by-BR matrix using the cbind function. The distribution of each BR’s SSC-A values was then calculated by creating 512 equally sized SSC-A value bins, based on the minimum and maximum recorded SSC-A values across all BR’s and samples, and recording the number of events greater than or equal to the bin value and less than the next highest bin value for each bin. To account for differences in the number of recorded events per BR, each BR’s SSC-A bin count values were normalized to the maximum count value recorded for that BR. Distributions were visualized as the average normalized count per SSC-A bin with standard deviation ribbons calculated per SSC-A bin and plotted as +/− the average per bin. Distributions were compared statistically using Satterthwaite’s method nested t-test.

### Migration and invasion assay

#### Low throughput

Melanocytes or melanoma cells were dissociated as described above and were plated at 150,000 cell/cm^2^ in a PO/Fibronectin/Laminin coated plate in MIM or tc-treated plate in DMEM 10%FBS, respectively. After 2 hours to allow cells to attach, wells were scratched with a 10ul pipette tip, using a new tip for each well. After scratching, a media change was performed to remove any free floating and/or dead cells and, optionally, to start drug treatments. This media change was considered time 0hr of the assay and representative brightfield images were taken on an echo revolve microscope to establish the average initial scratch width. Scratched plates were cultured for 72hrs and then cells were washed with PBS. After washing, cells were fixed with 4% PFA for 30min at room temperature. After fixation cells were washed with PBS and then incubated in 5ug/ml WGA, Alexa Fluor® 647 (Thermo) conjugate for 10min at room temperature. After incubation with WGA, cells were washed 3x with PBS and then left in PBS to image on an epi-fluorescent echo revolve microscope. Images were processed and analyzed using FIJI. Briefly, brightness and contrast was adjusted to maximize the cell membrane staining signal to background ratio. Next, images were thresholded to include cell membrane staining and exclude background. Then a region of interest (ROI) was drawn around the estimated original scratch area using the polygon tool, ensuring that the width of the ROI was equal to the width of the representative scratch images taken at 0hr and centered along the remaining 72hr scratch area. Migration efficiency into the scratched areas was quantified as the percentage of area covered by cells in the ROI. Statistical comparisons were performed using an unpaired t-test when comparing two sample groups and using a one-way ANOVA with multiple comparisons when comparing more than two sample groups.

#### High throughput

Melanocytes or melanoma cells were dissociated as described above and were plated at 150,000 cell/cm^2^ in Geltrex coated plates in MIM or DMEM 10% FBS, respectively. After 2 hours to allow cells to attach, wells were scratched with an AccuWound 96 Scratch Tool (Agilent). If scratching multiple plates at once, the scratch tool was washed between plates according to the supplier’s protocol. After scratching, a media change was performed to remove any free floating and/or dead cells and, optionally, add drug treatments. This media change was considered time 0hr of the assay and brightfield images were taken to establish the average initial scratch area in each well using a molecular devices ImageXpress high-content imaging system. Scratched plates were cultured for 72hrs and then cells were washed with PBS. After washing, cells were fixed with 4% PFA for 30min at room temperature. After fixation cells were washed with PBS and then incubated in 5ug/ml WGA, Alexa Fluor® 647 conjugate for 10min at room temperature. After incubation with WGA, cells were washed 3 times with PBS and then left in PBS to image on a molecular devices ImageXpress high-content imaging system. Images were analyzed using FIJI as described above.

### Doubling Rate Assay

Melanocytes or melanoma cells were dissociated with Accutase as described above and counted using a hemocytometer. Melanocytes were plated at 150,000 cell/cm^2^ in a PO/Fibronectin/Laminin coated plate in MIM. After allowing the melanocytes to attach for two hours, three wells were lysed with 1X RIPA buffer on ice for 5 minutes. The RIPA solution was pipetted up and down repeatedly in the well to lyse residual cells and then transferred to a microcentrifuge tube. Cell lysates were centrifuged at 14,000xg for 15 minutes at 4C. The supernatant was then transferred to a new microcentrifuge tube and diluted 1:50 in PBS and mixed with propidium iodide (PI) in PBS at a final ratio of 1:3000 before transferring to a 96 well plate. PI fluorescence intensity was measured by excitation at 535nm and emission detection at 615nm using a plate reader. The mean of the three wells was calculated to quantify the average DNA concentration per well at time 0hr. The remaining wells were cultured for 72hrs before repeating the RIPA lysis and PI based DNA concentration quantification steps. The growth rate of each well was calculated as (72hr PI a.u. -average 0hr PI a.u.)/average 0hr PI a.u. where the 0hr average was specific to each independent assay and sample. Sample groups were statistically compared using an unpaired t-test.

### Mice

C57BL6/J males and females were mated, where the presence of a vaginal plug was identified as E0.5. Pregnant female mice were euthanized at E12.5 and embryos were dissected. All mouse work was performed under the University of California, San Francisco, Institutional Animal Care and Use Committee guidelines in an approved facility of the Association for Assessment and Accreditation of Laboratory Animal Care International.

### Preparation of embryos

E12.5 embryos were dissected in 0.4 % BSA in PBS. Embryos were fixed rocking in 4% PFA overnight at 4°C. After fixation, embryos were washed three times with PBS for 20min each then cryoprotected by immersion in a 10% - 30% stepwise sucrose gradient overnight at 4°C. Embryos were incubated in 1:1 30% sucrose:OCT (Fisher) for 1hr, then embedded transversely in OCT for storage at −80 °C. Embryos were sectioned (20 mm thickness) using a cryostat (Leica).

### Mouse Tissue quantifications

Tissue section tile scans were analyzed using FIJI. Max intensity projections were created and each channel’s brightness was manually adjusted. The multi-point tool was used to mark CD44+/− MITF+ nuclei ROIs. MITF and CD44 positivity was determined manually by signal relative to local background. Dorsoventral location of each MITF nuclei was calculated via measurement of the X and Y coordinates of the multipoint ROIs and then scaled to the X and Y coordinates of the most dorsal and most ventral point of each tissue section.

### Mouse tissue immunofluorescence staining and imaging

Transverse embryo sections from cranial, front limb, abdomen, and hind limb regions were washed in PBS followed by permeabilization with 0.4% Triton-X in PBS for 10min. Tissue sections were blocked with 5% donkey serum and % BSA in 0.1 % Triton-X in PBS for 2hr at room temperature. Tissue sections were incubated with MITF, TUBB3, CD44-AF647 primary antibodies diluted (Table 1) in blocking buffer overnight at RT. Antibodies were detected by incubating with secondary antibodies diluted 1:200 in blocking buffer overnight at RT. Samples were tile-imaged by using a white-light Leica TCS SP8 converted confocal microscope with a 25 X water objective, 0.75 X optic zoom, and 1024 x 1024 pixel resolution, and stacks were acquired at system-optimized z steps between optical sections (z step size, 1 um).

### Generation of MeWo and A375 CRISPRi lines

A dCas9-KRAB-mCherry lentiviral construct (pHR-UCOE-EF1A-dCas9-HA-2xNLS-XTEN80-KRAB-P2A-mCherry)^42^ was used to generate CRISPRi-competent MeWo and A375 cells. HEK293T cells were maintained in DMEM supplemented with 10% FBS, penicillin, streptomycin and amphotericin B, and passaged using 0.05% Trypsin prior to cells reaching 90% confluency. To generate dCas9-KRAB-mCherry lentivirus for transduction of the melanoma cell lines, HEK293T cell were first plated at 175,000 cells/cm2. The dCas9-KRAB-mCherry plasmid was mixed with the pCMV_ΔR8.91 and pMD2.G packaging vectors at a ratio of 9:8:1 with TransIT®-Lenti Transfection Reagent (Mirus) in optiMEM (Gibco) and used for the HEK293T cell transfection. After 3 days, lentivirus-containing media was harvested and filtered using a .45um filter. The viral supernatant and polybrene (8 ug/ml final concentration, Sigma) was then added to the melanoma cell lines, both of which already contained luciferase reporter constructs for *in vivo* imaging, plated at 500,000 cells per 10cm plate. Transduced cell lines were then expanded and purified using fluorescence activated cell sorting for the mCherry reporter. CRIPSRi activity in each cell line was validated by lentiviral transduction with pCRISPRia-v2 (Addgene #84832) backbones containing either a guide RNA (gRNA) targeting ST3GAL4 or a non-targeting gRNA (Table 5) using the above protocol. Following transduction with the guide expression constructs, transduced cells were purified using puromycin (Gibco) selection. Once purified, RNA was extracted using the PureLink RNA kit and knockdown was assessed by qPCR as described above using primers to detect ST3GAL4 expression (Table 5). The line was determined efficient if ST3GAL4 expression was reduced by 80%.

### gRNA library generation

The gRNA library was designed to contain five gRNAs targeting each transcriptional start site of the top 100 most significantly enriched genes in the T1 and T2 melanocytes when directly compared as well as 50 non-targeting control gRNAs for a total of 1125 gRNAs. gRNA sequences were obtained from the previously described human genome-wide CRISPRi (v2.1) gRNA library^58^ and inserted between flanking nucleotide sequences to be used for PCR amplification and ordered as an oPool from IDT (Table 5). The PCR-amplified gRNA pool was then cloned into the pCRISPRia-v2 backbone via the BstXI-Bpu1102I (Thermo) sites and electroporated into competent *E. coli* cells (Thermo). Colony-forming unit (CFU) quantification of the electroporated library was used to assess library coverage (>5000x CFU per library element coverage achieved). The gRNA sequences were PCR-amplified from the library plasmid pool and sequenced on the Illumina MiSeq sequencer to confirm the uniform gRNA representation and distribution.

### CRISPRi i*n vivo* screen

gRNA library lentivirus was generated as described above, aliquoted and frozen. Viral titer was approximated by infection of MeWo cells with titrated virus dilutions followed by flow cytometry, determining the percentage of BFP-expressing cells. For the screen, 3 million MeWo cells per replicate were transduced with gRNA library virus at 30% MOI to ensure >500x library coverage in the transduced population. Transduced cultures were puromycin selected and immediately used for the following steps to avoid fitness bias. First, two replicates were collected immediately for genomic DNA extraction to serve as reference for the starting gRNA pool at time 0. Two additional replicates were tail vein injected into NSG (NOD scid gamma, Jackson Labs) mice. Lung colonization was monitored by *in vivo* luciferase activity imaging for 6.5 weeks, the animals were euthanized, lung tissue was collected, roughly mechanically dissociated and fully dissociated by incubation in 1mg/ml liberase (Sigma) in 1:1 DMEM:F12K supplemented with DNase I (Worthington Biochemical), 6.5mg/ml collagenase (Worthington Biochemical), penicillin, streptomycin and amphotericin B at 37°C with shaking. If necessary, red blood cell lysis was performed with ACK buffer (Thermo) before mouse cells were removed from the dissociated lungs using the mouse cell depletion kit (Miltenyi Biotec) yielding purified MeWo cells that achieved successful lung colonization. In parallel, two replicate cell populations were continued in culture *in vitro* for 17 days to mimic the estimated number of doublings achieved by the injected MeWo cells before genomic DNA extraction. Genomic DNA extraction for all samples was performed with QuickDNA (Zymo) kits. Each genomic DNA sample was digested with BstXI, and tagged by UMI-introducing primer extension (Table 5) using Klenow polymerase, exo-(NEB). The tagged gRNA sequences were PCR-amplified and sequenced on Illumina HiSeq 4000 sequencer SE50 run at UCSF Center for Advanced Technology. gRNA sequencing data was analyzed using iAnalyzeR (https://github.com/goodarzilab/iAnalyzer). Briefly, count matrices per gRNA were generated for each sample by alignment to the input gRNA protospacer sequences. gRNA abundances per sample were compared using a univariate analysis. Hits were selected as guides significantly decreased in metastatic samples compared to the input samples but not significantly decreased in *in vitro* samples compared to the input samples.

### Generation of MeWo and A375 knockdown cell lines

For each gene (HMG20B, MMP17 and SNRPB), two DNA oligos consisting of the top two predicted targeting protospacer sequences from previously described human genome-wide CRISPRi (v2.1) gRNA library^58^ were synthesized (IDT) and Gibson-cloned into pCRISPRia-v2 vector (Table 5). Both the MeWo and A375 CRIPSRi/Luciferase cells were transduced with gRNA-expressing lentivirus, produced in HEK293T cells as described above. Each gRNA knockdown cell line was puromycin selected and the knockdown efficiency was validated by qPCR.

### Bulk RNA sequencing and spliceoform analysis

RNA from control and SNRPB-KD MeWo and A375 samples was collected and isolated as described above. RNA-seq libraries were prepared using SMARTer Stranded Total RNA-Seq Kit v3 - Pico Input Mammalian (Takara) according to manufacturer’s recommendations, using 10 ng total RNA as input. The libraries were sequenced on a 2x 35 bp PE run on a NextSeq 550 instrument (Illumina). UMI-tools (v1.0.0) was used to extract UMI’s from read 2. Cutadapt (v2.3) was used to trim R1 to an identical length. This is required since MISO expects reads of equal length for both pairs. STAR (v2.7.1a) was then used to align reads to the human genome (hg38) and UMI collapse (1.0.0) was used to remove PCR duplicates (with --two-pass --paired --remove- unpaired --remove-chimeric flags). The resulting deduplicated bam files were sorted by name and converted back to fastq files (samtools v1.7). The deduplicated fastq files were aligned to a curated human transcriptome using bowtie (2.3.5) and bamPEFragmentSize (3.5.1) was used to calculate fragment length distribution parameters (i.e. mean and standard deviation). MISO (0.5.4) was then used to analyze changes in skipped exon inclusions by passing the deduplicated STAR-aligned bam files along with the fragment size distribution for each sample.

### SNRPB CLIP

HEK-293 cells with SNRPB endogenously tagged with split mNeonGreen (Cho et al. Science 2022) were crosslinked with 400mJ/cm2 254nm UV. Crosslinked cells were lysed with ice-cold lysis buffer (1X PBS, 0.1% SDS, 0.5% sodium deoxycholate, 0.5% IGEPAL CA-630) supplemented with 1x halt protease inhibitors (Pierce) and SuperaseIN (Invitrogen). After lysis, samples were treated with DNase I (Promega) at 37degC, shaking at 1000rpm for 5 minutes. Each lysate sample was then split equally, and each portion treated with either a medium or low dilution of a mix of RNase A and RNase I (Thermo; 0.11ng/ul RNase A and 0.03units/ul RNase I, and 0.02 RNase A and 0.03units/ul RNase I, respectively) at 37degC for 5 minutes. Lysate was cleared by centrifuging at 21,000xg 4degC 20 minutes. The clarified lysate was added to pre-washed mNeonGreen-trap magnetic agarose beads (ChromoTek). Immunoprecipitation was performed at 4degC with end-over-end rotation for 2 hours. Beads were then washed 2X with high salt wash buffer (5X PBS, 0.1% SDS, 0.5% sodium deoxycholate, 0.5% IGEPAL CA-630), 2X with lysis buffer (1X PBS, 0.1% SDS, 0.5% sodium deoxycholate, 0.5% IGEPAL CA-630), and 2X with PNK buffer (10mM Tris-HCl pH 7.5, 0.5% IGEPAL CA-630, 10mM MgCl2). The RNA was then end-repaired and poly(A) tailed on-bead by treatment first with T4 PNK (NEB) and then treatment with yeast poly(A) polymerase (Jena Bioscience). The RNA was further 3’-end-labeled with azide-dUTP (TriLink biotechnologies) using yeast poly(A) polymerase (Jena Bioscience). The protein-RNA complexes were then labelled with IRDye800-DBCO (LiCor). The protein-RNA complexes were then eluted from beads, resolved by running on a 4-12% Bis-Tris NuPAGE gel (Invitrogen), transferred to protran BA85 nitrocellulose (Cytiva), and imaged using an Odyssey Fc instrument (LiCor). The regions of interest were excised from the membrane and the RNA was isolated by Proteinase K (Invitrogen) digestion and subsequent pulldown with oligodT dynabeads (Invitrogen). After eluting from the oligodT beads, the RNA was immediately used for library preparation using the SMARTer smRNA-Seq Kit (Takara), with the following modifications. The poly(A) tailing step was omitted, and reverse transcription was performed with a custom RT primer. The PCR step was performed with indexed forward (i5) primers and a universal reverse (i7) primer. The libraries were PAGE purified and then sequenced on an Illumina HiSeq4000 instrument at the UCSF Center for Advanced Technologies.

### Single cell RNA sequencing (scRNAseq)

#### Library preparation, sequencing and processing

Adherent or 3D cultures were dissociated using the methods described above specific to each timepoint collected. Single cell suspensions were counted and diluted for GEM generation & barcoding, post GEM-RT cleanup & cDNA amplification, 3’ gene expression library construction, and sequencing on Illumina NovaSeq sequencer according to the 10x Genomics Chromium Next GEM Single Cell 3’ Reagent Kits v2 User Guide. CellRanger v2.1.1 was used to map reads to the human reference hg38 transcriptome and generate gene count matrices.

### scRNAseq data analysis

#### Cell quality assessment and filtration

Each dataset, corresponding to a unique timepoint or differentiation protocol was analyzed independently using Seurat (v4.3). First, several per-cell quality control metrics were calculated such as the percentage of mitochondrial gene reads (percent.mito), the percentage of ribosomal gene reads (percent.ribo) and a gene per umi novelty score calculated as log10(nFeatures) divided by the log10(nCount)(novelty.score). Low quality cells were identified using nCount and nFeature ranges specific to each dataset (Table 2) and excluded from further analysis.

#### Clustering, dimensionality reduction and annotation

Datasets were log normalized using a scale factor of 10,000 using NormalizeData. Variable features were identified using the “vst” method with default parameters of the FindVariableFeatures function. Cell cycle status was predicted using the CellCycleScoring function based on Seurat provided S phase and G2M Phase gene sets. Feature counts were scaled and the calculated S.Score, G2M.Score, percent.ribo and novelty score variables were regressed out using ScaleData. Principal component analysis was performed using RunPCA with default parameters. Uniform manifold approximation and projection (UMAP) dimensionality reduction was performed using RunUMAP with PCA as the reduction input. Shared nearest-neighbor (sNN) graphs were constructed using FindNeighbors. Finally, cell clusters were unbiasedly identified using louvain clustering via the FindClusters function. The number of principal components used in RunUMAP and FindNeighbors as well as the resolution used in FindClusters was specific to each dataset (Table 3). Differentially expressed genes of each cluster were identified using the Wilcoxon Rank Sum test with the FindAllMarkers function. Clusters were annotated using known cell-type markers. Gene-dropout correction via count imputation was performed using adaptively-thresholded low rank approximation (ALRA) using the RunALRA function in the SeuratWrappers package (v.3). Imputed gene counts were used for all subsequent analysis unless otherwise specified^59,60^.

#### Transcription factor activity analysis

SCENIC (v1.3) was run independently on each dataset. Imputed count matrices and metadata were extracted from the above analyzed Seurat objects. Human RcisTarget databases were downloaded from resources.aertslab.org. Imputed count matrices filtered to remove genes with total counts less than 3% of the number of cells in the dataset or cells with detected genes fewer than 1% of the number of cells in the dataset. Coexpression networks were identified using runCorrelation and GENIE3 (v1.18) was used to identify transcription factor modules using runGenie3 from the filtered imputed matrix. Gene regulatory networks were then identified and scored using the SCENIC wrapper functions with default parameters. Briefly, runSCENIC_1_coexNetwork2modules was used to create the co expression modules, runSCENIC_2_createRegulons was used to identify regulons using transcription factor motif analysis and finally, regulon activity was calculated using runSCENIC_3_scoreCells.

#### Pseudotime analysis

Monocle3 (v1) was run independently on each dataset. Imputed count matrices and metadata were extracted from the above analyzed Seurat objects and used to create CellDataSets compatible with monocle3 using new_cell_data_set. The CellDataSet was preprocessed but not normalized, as the imputed counts matrix was already normalized, using preprocess_cds with norm_method = “none”. The number of dimensions used for each dataset during pre-processing was specific to each dataset and can be found in (Table 2). Cell cycle effects were removed using the align_cds function based on the cell cycle phases determined by Seurat CellCycleScoring. UMAP dimensionality reduction was performed using reduce_dimension. Clustering was not performed as cell cluster labels were transferred from the Seurat object. Finally, the pseudotime trajectories were predicted using the learn_graph function. A root state was chosen using order_cells based on the most central root point in the unbiased NCC cluster.

#### Ligand-receptor interaction analysis

CellChat (v1.4) was run independently on each dataset.Imputed count matrices and metadata were extracted from the above analyzed Seurat objects and used to create CellChat objects using createCellChat. All cell-cell communication types were used from the human CellChat database. The subsetData function was used to filter the imputed counts matrix for signaling related genes. Using the filtered maxtrix, overexpressed ligand and receptors are then identified per each specified cell grouping using the identifyOverExpressedGenes functions followed by over expressed ligand receptor interactions using identifyOverExpressedInteractions. The probability of communication between the specified cell groups is then predicted using the computeCommunProb function. The communication probability of whole signaling pathways were then computed based on the individual receptor-ligand probabilities using computeCommunProbPathway.

#### Module scoring

To identify transcriptionally similar cell populations between datasets, cluster gene signatures were created. First, the differentially expressed genes with a positive log2 fold change for each cluster in the reference dataset were filtered to exclude genes not detected in the query dataset. Of the remaining genes, the top 100 most significantly differentially expressed were used as the cluster’s transcriptional signature (Table 4). The query dataset’s cells were then scored for the expression of the cluster’s transcriptional signature using Seurat’s AddModuleScore function based on the imputed gene counts.

#### Similarly weighted non-negative embedding (SWNE) analysis

For the datasets to be compared, the counts matrices were filtered to only include genes detected in both the query and reference dataset. First, the similarly weighted nonnegative embeddings (SWNE) are calculated for the reference dataset using the SWNE (v0.6) package. Briefly, 3000 variable genes are identified from the reference dataset’s non-imputed gene counts using FindVariableFeatures and component factors are identified by nonnegative matrix factorization (NMF) using RunNMF from NMF v0.23. The number of factors to be identified (k) was set equal to the number of PCs used during initial analysis with Seurat) (Table 2). sNN graphs calculated previously were extracted from the reference seurat object and pruned using PruneSNN based on the reference dataset’s previously calculated kNN matrix, specified cell grouping and a Q-value cutoff of 0.001. Finally, Sammon mapping is used for dimensionality reduction of the component factors by calculating factor embeddings using the reference dataset’s sNN matrix. This allows for both cells and genes to be plotted based on their influence by and contribution to the component factors, respectively. To project the query dataset onto the reference SWNE, sNN between the query and reference dataset are first calculated. To do this, the reference dataset’s PC loadings are used to calculate PC embeddings of the previously determined 3000 variable gene based on the query dataset’s scaled gene counts. The sNN matrix is then calculated from the query dataset’s PC embeddings trained on the reference PC embeddings using ProjectSNN. Next, NMF gene loadings for the query dataset are calculated from the reference dataset’s factor decomposition results using ProjectSamples. Finally, the query dataset is projected onto the reference SWNE embeddings using ProjectSWNE.

#### Transcription factor modules identification

For Stage 2-4 NCC subset datasets, average regulon activity scores per NC subtype were calculated and the subsequent regulon by NC subtype matrices were merged, excluding S4 NCC3 which is not predicted to follow a melanogenic lineage. Regulon modules were identified using specific filtering criteria to identify regulons exhibiting desired trends in activity along the predicted lineage paths. Module 1 consists of regulons detected to be active in all subtypes, where the activity in S2 NCC2 is greater than S2 NCC1, S3 NCC2 is greater than S3 NCC3 by at least 0.05, and S3 NCC2 is greater than S3 NCC1 and all S2 and S4 NCC subtypes. Modules 2 consists of regulons detected to be active only in S2 and S3 subtypes, where the activity in S2 NCC2 is greater than S2 NCC1, S3 NCC2 is greater than S3 NCC3 by at least 0.05, and S3 NCC2 is greater than all S2 subtypes and S3 NCC1. Module 3 consists of regulons detected to only be active in S3 subtypes, where the activity of S3 NCC2 is greater than S3 NCC3 by at least 0.05, and S3 NCC2 is greater than S3 NCC1. Module 4 consists of regulons detected to only be active in S3 and S4 subtypes, where the activity of S3 NCC3 is greater than S3 NCC1, S3 NCC2 is greater than S3 NCC3 by at least 0.05, and S4 NCC1 is greater than S4 NCC3. Module 5 consists of regulons detected to be active in all subtypes, where the activity of S3 NCC3 is greater than S3 NCC2 and S4 NCC3 is greater than S2 NCC2 by at least 0.05. Module 6 consists of regulons detected to only be active in S3 and S4 subtypes, where the activity of S3 NCC3 is greater than both remaining S3 subtypes, and S4 NCCs is greater than S2 NCC2 by at least 0.05. Module 7 consists of regulons detected to be active only in S4 subtypes, where S4 NCC3 is greater than S4 NCC1 by at least 0.05. Finally, module 8 consists of regulons detected only S2 and S4 subtypes, where S4 NCC3 is greater than S4 NCC1 by at least 0.05 and S2 NCC2 is less than S2 NCC1 by at least 0.05.

#### Ligand-receptor module identification

For stage 2-4 NCC subset datasets, receptor-ligand signaling scores for each NC subtype were calculated and the subsequent regulon by NC subtype matrices were merged, excluding S4 NCC3 which is not predicted to follow a melanogenic lineage. Raw signaling scores were scaled by a factor of 10^6^ and log transformed for comparison and visualization. Ligand-receptor pair (LRP) modules were identified using specific filtering criteria to identify LRPs exhibiting desired trends in activity along the predicted lineage paths. Module 1 consists of LRPs active in S2 NCC1 and S3 NCC2, and the activity of S3 NCC2 is greater than S3 NCC3 by at least .7. Module 2 consists of LRPs active in S3 NCC1 and S3 NCC2, where the activity of S3 NCC2 is greater than S3 NCC3 by at least .7, and not active in S2 NCC1. Module 3 consists of LRPs active in S3 NCC2, where the activity of S3 NCC2 is greater than S3 NCC3 by at least .7, and not active in S2 NCC1 or S3 NCC1. Module 4 consists of LRPs active in S2 NCC1, S3 NCC3 and S4 NCC3, where both S3 NCC3 and S4 NCC3 are greater than S3 NCC2 by at least .7. Module 5 consists of LRPs active in S3 NCC1, S3 NCC3 and S4 NCC3, where both S3 NCC3 and S4 NCC3 are greater than S3 NCC2 by at least .7, but not active in S2 NCC1. Finally, Module 6 consists of LRPs active in S3 NCC3 and S4 NCC3, where both S3 NCC3 and S4 NCC3 are greater than S3 NCC2 by at least .7, but not active in S2 NCC1 or S3 NCC1.

### Analysis of published datasets

#### Mouse Neural Crest Dataset

QC filtered and processed .h5ad files were obtained from GEO (GSE201257). The reticulate package (v1.28) was used to interface with the python scanpy package to import data into R and was then converted into a Seurat object, preserving the authors original metadata, leiden clustering and UMAP coordinates. Leiden clusters were annotated into cell types using the expression of cell type markers described in the original publication^28^.

#### Human Melanocyte Datasets

Eight single cell RNA seq datasets containing melanocytes from healthy donors were curated from GEO (GSE151091, GSE162054/GSE153760, GSE147424, GSE150672 and GSE130973), the EGA (EGAS00001002927) and the human cell atlas: Developmental (https://developmental.cellatlas.io/) (Developmental cell programs are co-opted in inflammatory skin disease: Human Adult Healthy 10x Data and Human fetal 10x data). When necessary, datasets were first subsetted to exclude disease samples. All datasets were processed using the Seurat analysis pipeline described above with one modification. In place of running PCA analysis, mutual nearest neighbors (mnn) correction was performed using the FastMNN function from the SeuratWrappers package to perform batch correction of each unique patient sample within the respective dataset. Subsequently, the UMAP and sNN were calculated using the mnn reduction based on 30 dimensions. For datasets where uploaded metadata did not include cell type annotations, the melanocyte cluster was identified by expression of MITF, PMEL, DCT, TYR and MLANA. For each dataset the melanocyte cluster was subsetted and the relevant metadata names and levels were made consistent across datasets. All melanocyte datasets were then merged and processed using the same Seurat pipeline as above, utilizing the FastMNN function to perform batch correction based on the original dataset identifier.

### Analysis of datasets from the cancer genome atlas (TCGA)

BulkRNA sequencing datasets of 471 melanoma samples from 469 different patients were curated from the cancer genome atlas and merged into a sample by gene FPKM matrix. Ensembl IDs were converted to gene names using BiomaRt.

### Gene set enrichment analysis (GSEA)

Differential gene expression was performed comparing T1 to T2 melanocytes using the Wilcoxon Rank Sum test. T1 and T2 melanocyte transcriptional signatures were constructed consisting of the top 100 most significantly enriched genes in each respective melanocyte type. To identify melanoma samples that transcriptionally resemble either melanocyte type, gene set enrichment analysis was performed on each melanoma sample for the T1 and T2 melanocyte transcriptional signatures using the fgsea package (v1.22). Melanoma samples were labeled as either enriched or reduced for each transcriptional signature corresponding to a significant positive or significant negative enrichment score, respectively. Survival curves were generated using the patient metadata obtained from TCGA with the melanoma samples grouped by indicated melanocyte signature enrichment status and compared in Prism using the Mantel-Cox test. When comparing T1 enriched to T2 enriched melanomas, samples that were significantly enriched for both signatures were discarded from the analysis.

### Survival analysis

Survival analysis using the melanoma TCGA data was used to identify higher confidence hits from the CRISPRi screen. For all hits nominated by the univariate analysis, the full range of expression values was tested as thresholds for grouping samples with an expression value higher or lower than the test threshold and calculating the survival difference between the high and low expression ground using the survival package (v3.5). Genes were considered high confidence hits if an expression threshold for that gene existed where higher expressing samples had significantly lower survival probability than lower expressing samples.

### Cox proportional hazard analysis

To further narrow high confidence gene targets, a cox proportional hazard was performed for all genes with significant survival thresholds using coxph. Using the expression threshold determined in the survival analysis, both a univariate and multivariate cox proportional hazard were performed. Univariate tests considered only the samples higher or lower expression than the predetermined threshold, while the multivariate test also considered multiple other criteria from the patient metadata (diagnosis age, metastasis stage, lymph node stage, neoplasm disease stage, tumor stage, neoadjuvant therapy type, adjuvant postoperative pharmaceutical therapy, adjuvant postoperative radiotherapy and primary or metastatic sample type). Genes were considered high confidence hits if the expression threshold grouping maintained a significant hazard ratio when all variables were considered.

---

**Figure S1:**
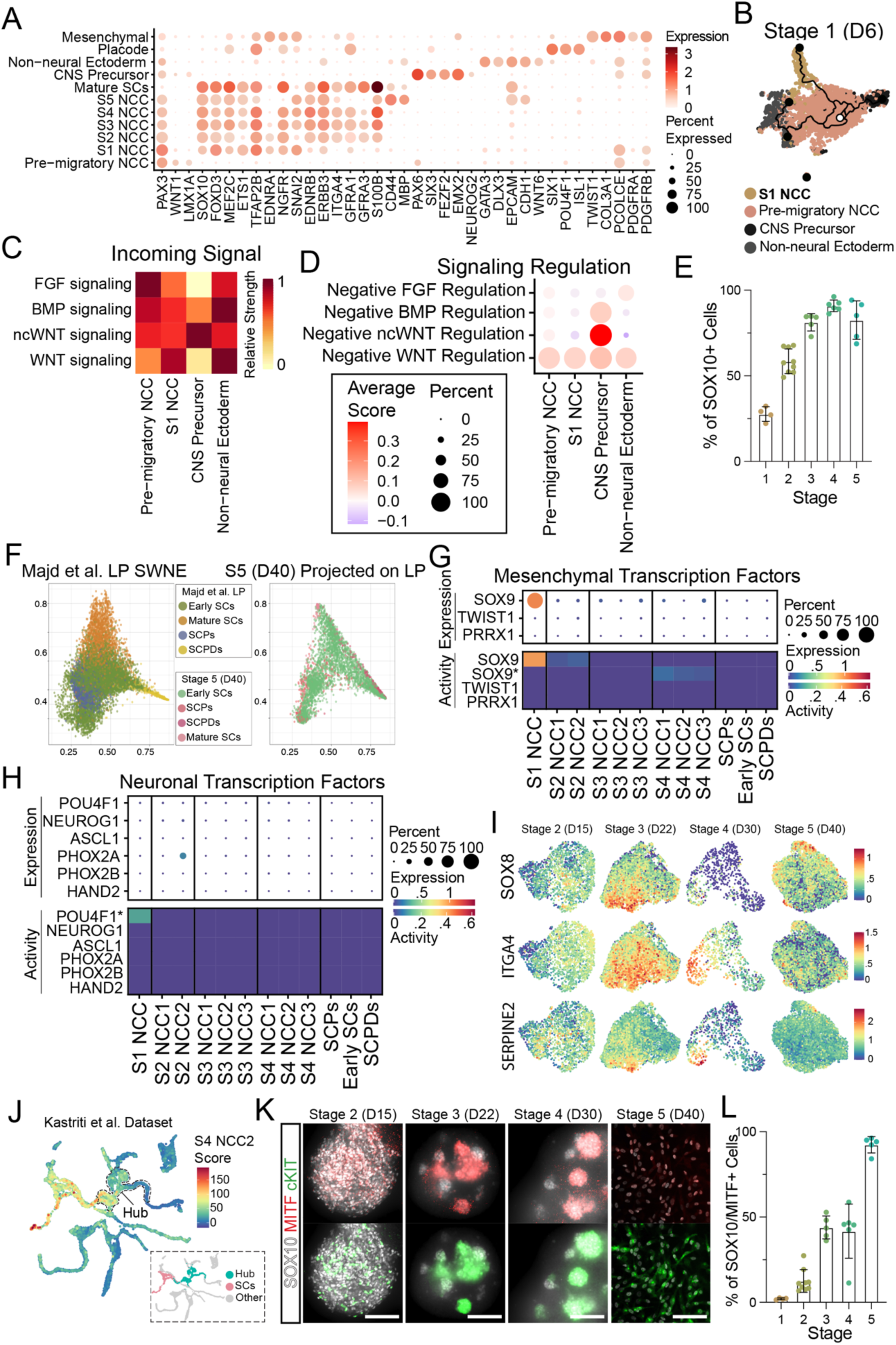
Differentiation of hPSCs toward NCC and Schwann cells yields heterogeneous populations with different lineage potentials. A. Expression of canonical cell type markers in stage 1-5 clusters used for cell type annotation. B. Pseudotime lineage prediction of stage 1 neural crest induction. Cells (points) are colored by cell type cluster. Black lines follow predicted paths of transcriptional change from selected starting nodes (white circle) to terminal nodes (black circle). C. Predicted relative strength of cell type response to incoming signaling pathways necessary for neural plate patterning. D. Modules scoring of gene ontology gene lists for negative regulation of neural plate patterning signals. E. Flow cytometry-based quantification of SOX10 positive populations in stage 1-5 crestospheres and stage 5 SCP induction. Error bars represent standard deviation. F. Projection of stage 5 Schwann subclusters and cell types into the SWNE embeddings of the “low passage” Schwann induction dataset published in Majd et al. Cells (points) with similar SWNE coordinates are transcriptionally similar to one another. Abbreviations: SCs = Schwann Cells, SCPs = Schwann Cell Precursors, SCPDs = Schwann Cell Precursor Derivatives. G. Expression (top) and predicted activity (bottom) of canonical mesenchymal transcription factors in stage 1-5 NCC and Schwann subclusters. Asterisks represent TF activity predicted from lower confidence motif annotations. H. Expression (top) and predicted activity (bottom) of canonical sensory and autonomic neural transcription factors in stage 1-5 NCC and Schwann subclusters. Asterisks represent TF activity predicted from lower confidence motif annotations. I. Feature plots of normalized expression for hub markers SOX8 (top), ITGA4 (middle) and SERPINE2 (bottom) in stage 2-5 NCC and Schwann subclusters. J. Feature plot Kastriti et al. dataset colored by module score for S4 NCC2 top 100 marker genes. Dotted line outlines the cells annotated as “Hub” by the original authors. UMAP inlay colored by author annotations for the Hub and SC cells. K. SOX10 (white), MITF (red) and cKIT (green) expression in stage 2-4 crestospheres and stage 5 SCP induction. Scale bar represents 100um. L. Flow cytometry-based quantification of SOX10 positive and SOX10/MITF double positive populations in stage 1-5 crestospheres and stage 5 SCP induction. Error bars represent standard deviation.

**Figure S2:**
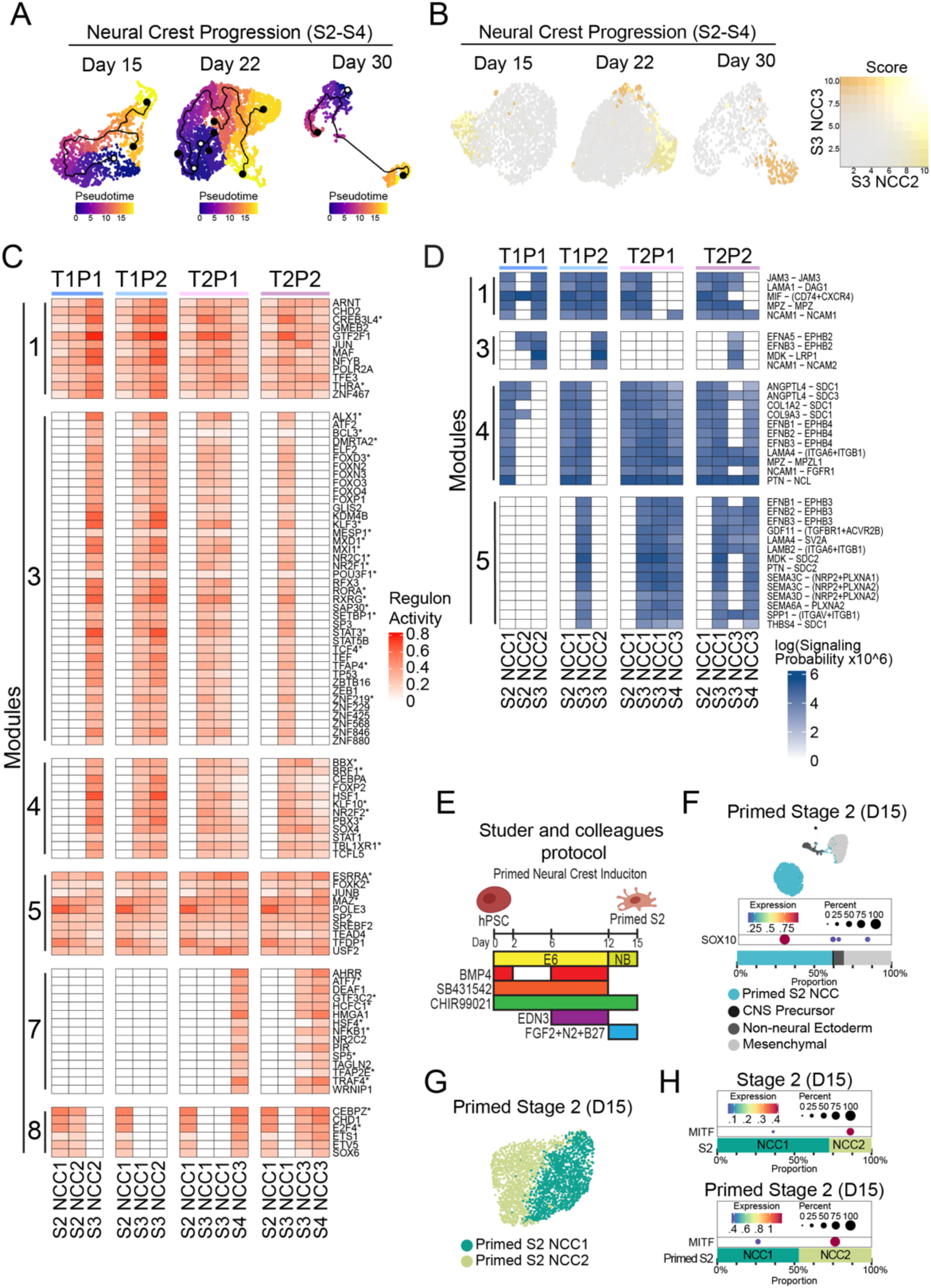
Distinct melanogenic trajectories show different transcriptional and signaling programs. A. Pseudotime lineage reconstruction of stage 2-4 neural crest subtypes. Cells (points) are colored by pseudotime value. Black lines follow predicted paths of transcriptional change from selected starting nodes (white circle) to terminal nodes (black circle). B. Feature plots of stage 2-4 NCCs colored by a blended module score for S3 NCC2 (yellow) and S3 NCC3 (orange) respective top 100 marker genes. C. Heatmap of predicted transcription factor activity per stage 2-4 NC subtypes for transcription factor modules 1, 3, 4, 5, 7, and 8. NC subtypes are grouped by proposed lineage path progressions. D. Heatmap of predicted ligand receptor pair activity per stage 2-4 NC subtypes for ligand receptor pair modules 1, 3, 4, and 4. NC subtypes are grouped by proposed lineage path progressions. E. Schematic of directed differentiation protocol published by Studer and colleagues to differentiate human pluripotent stem cells to melanogenic primed stage 2 neural crest cells with defined media conditions. The protocol differs from the previously described protocol due to the addition of BMP4 and EDN3 during days 6-12 of neural crest induction. F. UMAP embedding, SOX10 expression, cluster proportions, and cluster annotation of the primed stage 2 neural crest induction. G. UMAP embeddings and subclustering of the primed stage 2 SOX10+ cluster. H. MITF expression and cluster proportions of stage 2 and primed stage 2 NC subtypes.

**Figure S3:**
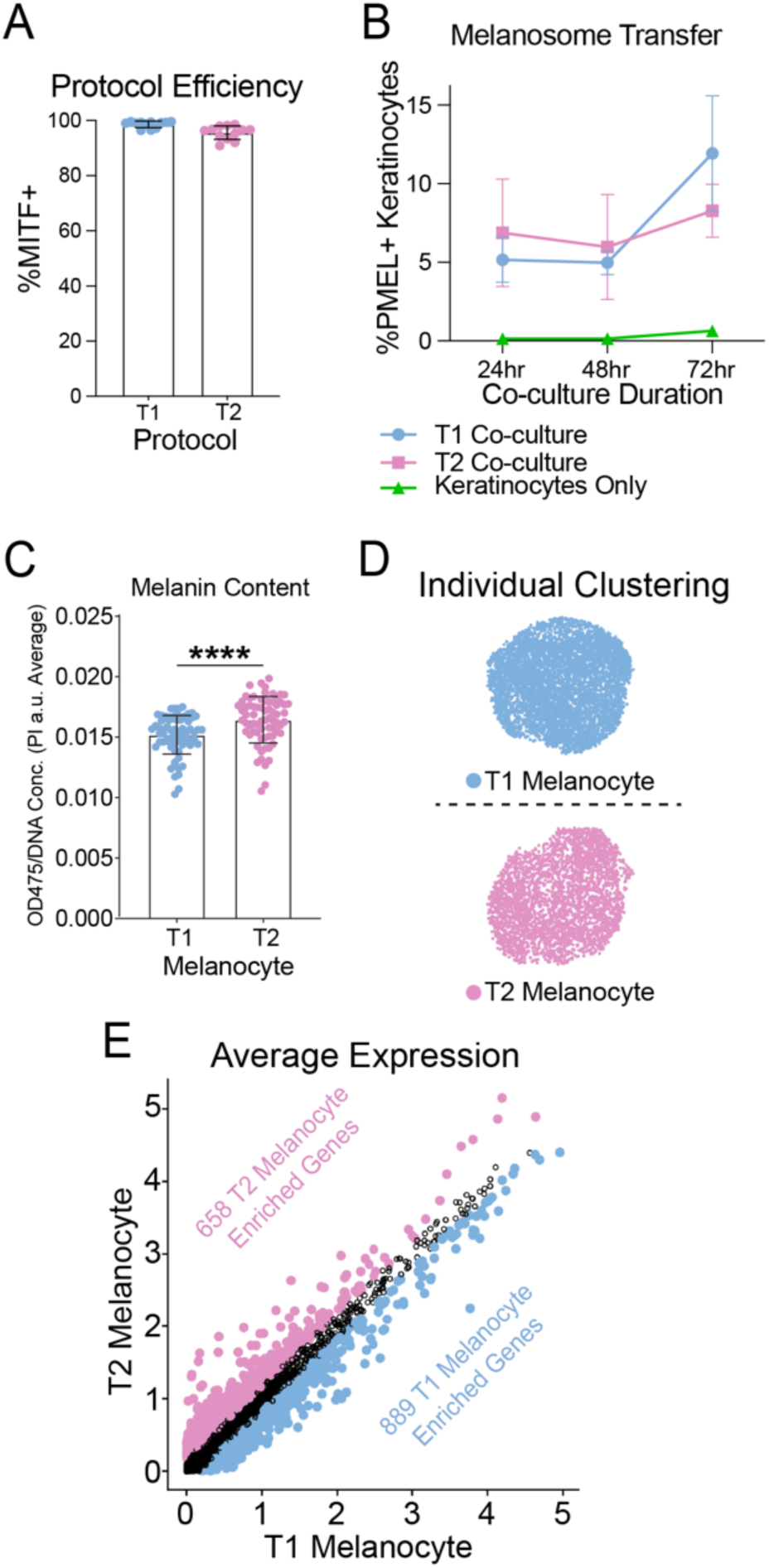
T1 and T2 melanocytes have different transcriptional signatures. A. Flow cytometry-based quantification of MITF+ cells resulting from the T1 and T2 melanocyte induction protocols (n=12). Error bars represent standard deviation. B. Flow cytometry-based quantification of melanosome transfer from T1 or T2 melanocytes to primary keratinocytes after 24, 48 and 72hrs of co-culture or with keratinocytes alone. The y axis represents the percentage of K14+/PMEL+ keratinocytes. Points represent technical replicates from n=3. Error bars represent standard deviation. C. Bulk culture pigmentation quantification based on optical density readings at 475nm normalized to DNA concentration based on propidium iodide fluorescence intensity. Points represent technical replicates from n=3. **** is p<0.0001 and error bars represent standard deviation. D. UMAP embeddings of individually analyzed T1 (top) and T2 (bottom) melanocyte datasets. E. Scatter plot of average expression of all detected genes in T1 and T2 melanocytes. Genes are colored by differential expression where genes in blue are significantly (p < 0.05) more highly expressed in T1 melanocytes by > 0.25 fold change while genes in pink are significantly (p < 0.05) more highly expressed in T2 melanocytes by > 0.25 fold change. Genes represented by black outlines are not significantly differentially expressed.

**Figure S4:**
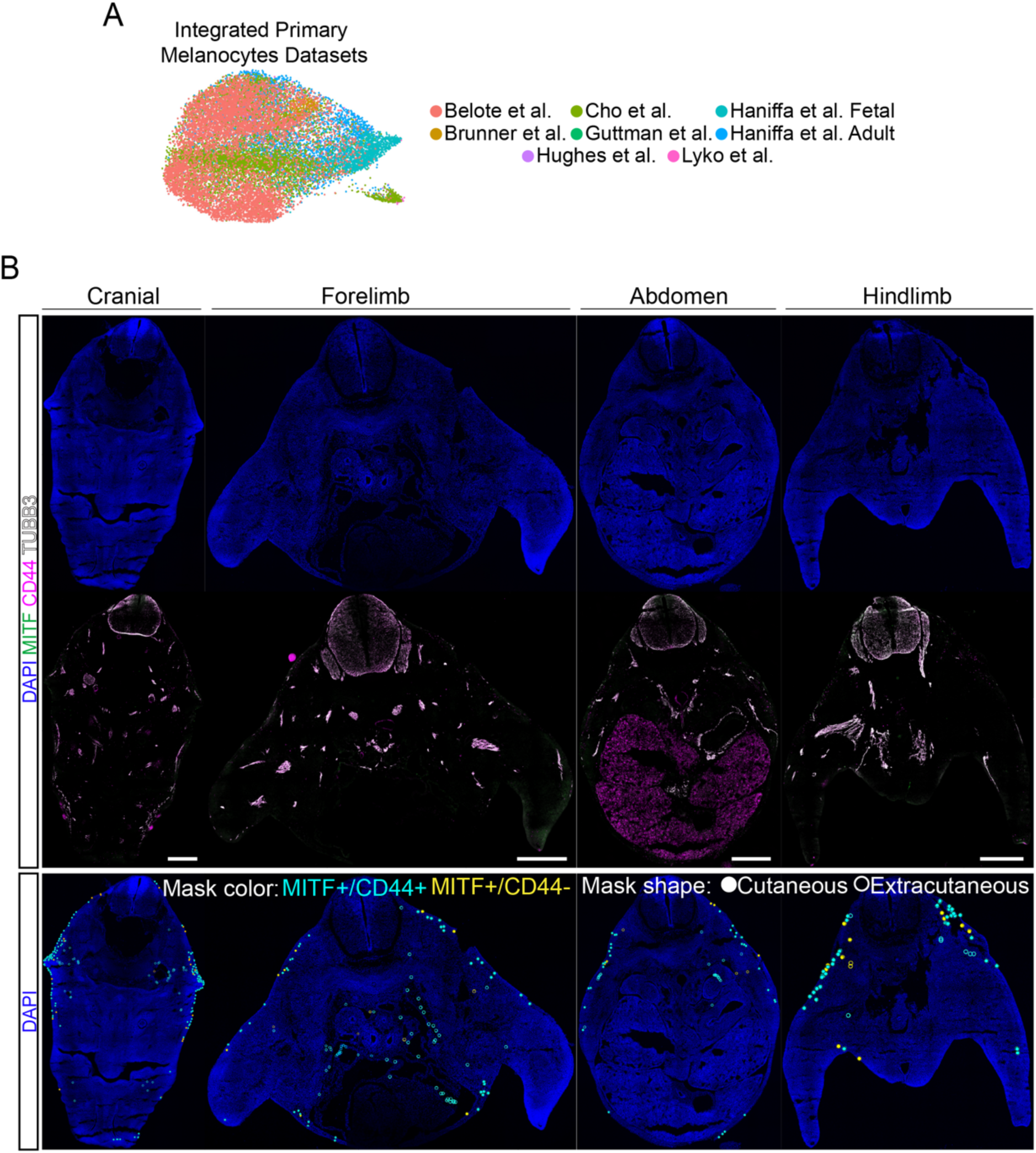
Evaluation of T1 and T2 markers in human adult and mouse fetal primary melanocytes. A. UMAP embedding of eight integrated primary human melanocyte datasets colored by the dataset’s original publication identifier. B. Top panel: Representative immunofluorescent images of E12.5 mouse cranial, front limb, abdominal and hind limb sections showing nuclei (blue), melanocytes (MITF, green) neurons (TUBB3, grey) and the T2 melanocyte surface marker CD44 (magenta). Bottom panel: Representative E12.5 mouse cranial, front limb, abdominal and hind limb sections showing nuclei (blue) and region of interest masks marking all observed MITF+ cells. Masks are colored based on CD44 expression (CD44+ = cyan, CD44-= yellow) and shaped based on anatomical position (closed circle = cutaneous, circle outline = extracutaneous). Scale bars represent 500um.

**Figure S5:**
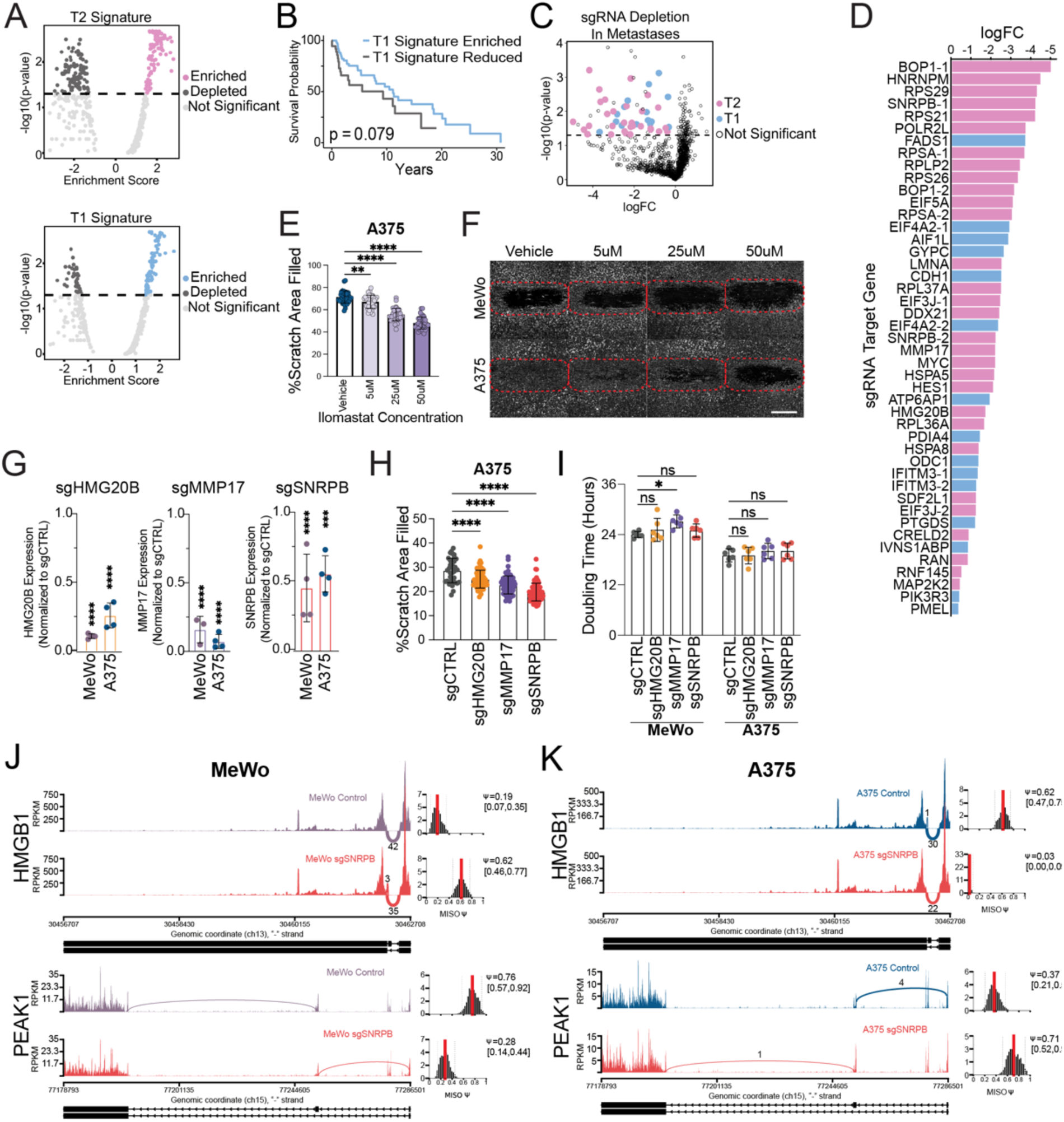
Functional roles of T1 and T2 melanocyte markers in melanoma. A. Gene set enrichment analysis of 471 Bulk RNAseq melanoma samples for the respective top 100 T2 (top) and T1 (bottom) marker genes. Melanoma samples are colored by their significant enrichment (pink/blue) or reduction (dark gray) for the gene sets, or light gray for non-significant. B. Survival curves of melanoma patients with melanomas significantly enriched (blue) versus significantly reduced (grey) for the expression of the top 100 T1 melanocyte markers. C. Volcano plot of gRNA enrichment or depletion in sequenced lung metastases compared to the culture prior to injection. gRNAs labeled in color are significantly depleted in the lung metastases but not significantly depleted the *in vitro* cultured pool. D. Bar chart of fold change of significantly depleted gRNAs in lung metastasis compared to starting abundance. Bars are colored by gRNA target of a T1 marker gene (blue) or T2 marker gene (pink). E. Scratch assay quantification of A375 cells treated with increasing concentrations of the MMP inhibitor ilamostat. Points represent technical replicates from n=3. p-values are: ** p<0.01, **** p<0.0001 and error bars represent standard deviation. F. Representative images of scratch assays of MeWo (top) and A375 (bottom) cells treated with increasing concentrations of the MMP inhibitor ilamostat. Scale bar represents 1000um. G. qPCR determined fold change expression of CRISPRi hit genes in stable MeWo and A375 KD lines after CRISPRi targeting normalized to non-targeting gene expression. p-values are: *** p<0.001, **** p<0.0001 and error bars represent standard deviation. H. Scratch assay quantification of CRIPSRi A375 cells transduced with gRNAs targeting the nominated genes versus a non-targeting control. Points represent technical replicates from n=3 assay replicated with n=2 gRNAs. **** p<0.0001 and error bars represent standard deviation. I. Doubling rate analysis of CRISPRi MeWo and A375 cells transduced with gRNAs targeting the nominated genes versus a non-targeting control. Points represent technical 2 replicates from n=3. p-values are: * p<0.05, ns not significant and error bars represent standard deviation. J. Sashimi plots showing the differential splicing patterns of migration related genes HMGB1 and PEAK1 in non-targeted versus SNRPB KD MeWO melanoma cell lines. K. Sashimi plots showing the differential splicing patterns of migration related genes HMGB1 and PEAK1 in non-targeted versus SNRPB KD A375 melanoma cell lines.

## References

1. Bronner, M.E., and LeDouarin, N.M. (2012). Development and evolution of the neural crest: an overview. Dev. Biol. 366, 2–9. 10.1016/j.ydbio.2011.12.042.

2. Dupin, E., Real, C., Glavieux-Pardanaud, C., Vaigot, P., and Douarin, N.M.L. (2003). Reversal of developmental restrictions in neural crest lineages: Transition from Schwann cells to glial-melanocytic precursors in vitro. Proc. Natl. Acad. Sci. 100, 5229–5233. 10.1073/pnas.0831229100.

3. Baggiolini, A., Varum, S., Mateos, J.M., Bettosini, D., John, N., Bonalli, M., Ziegler, U., Dimou, L., Clevers, H., Furrer, R., et al. (2015). Premigratory and migratory neural crest cells are multipotent in vivo. Cell Stem Cell 16, 314–322. 10.1016/j.stem.2015.02.017.

4. Bronner-Fraser, M., and Fraser, S.E. (1988). Cell lineage analysis reveals multipotency of some avian neural crest cells. Nature 335, 161–164. 10.1038/335161a0.

5. Bronner-Fraser, M., and Fraser, S. (1989). Developmental potential of avian trunk neural crest cells in situ. Neuron 3, 755–766. 10.1016/0896-6273(89)90244-4.

6. McKinney, M.C., Fukatsu, K., Morrison, J., McLennan, R., Bronner, M.E., and Kulesa, P.M. (2013). Evidence for dynamic rearrangements but lack of fate or position restrictions in premigratory avian trunk neural crest. Dev. Camb. Engl. 140, 820–830. 10.1242/dev.083725.

7. Serbedzija, G.N., Fraser, S.E., and Bronner-Fraser, M. (1990). Pathways of trunk neural crest cell migration in the mouse embryo as revealed by vital dye labelling. Dev. Camb. Engl. 108, 605–612. 10.1242/dev.108.4.605.

8. Baroffio, A., Dupin, E., and Le Douarin, N.M. (1988). Clone-forming ability and differentiation potential of migratory neural crest cells. Proc. Natl. Acad. Sci. U. S. A. 85, 5325–5329. 10.1073/pnas.85.14.5325.

9. Stemple, D.L., and Anderson, D.J. (1992). Isolation of a stem cell for neurons and glia from the mammalian neural crest. Cell 71, 973–985. 10.1016/0092-8674(92)90393-q.

10. Krispin, S., Nitzan, E., and Kalcheim, C. (2010). The dorsal neural tube: A dynamic setting for cell fate decisions. Dev. Neurobiol. 70, 796–812. 10.1002/dneu.20826.

11. Harris, M.L., and Erickson, C.A. (2007). Lineage specification in neural crest cell pathfinding. Dev. Dyn. Off. Publ. Am. Assoc. Anat. 236, 1–19. 10.1002/dvdy.20919.

12. Henion, P.D., and Weston, J.A. (1997). Timing and pattern of cell fate restrictions in the neural crest lineage. Dev. Camb. Engl. 124, 4351–4359. 10.1242/dev.124.21.4351.

13. Fattahi, F., Steinbeck, J.A., Kriks, S., Tchieu, J., Zimmer, B., Kishinevsky, S., Zeltner, N., Mica, Y., El-Nachef, W., Zhao, H., et al. (2016). Deriving human ENS lineages for cell therapy and drug discovery in Hirschsprung disease. Nature 531, 105–109. 10.1038/nature16951.

14. Hackland, J.O.S., Shelar, P.B., Sandhu, N., Prasad, M.S., Charney, R.M., Gomez, G.A., Frith, T.J.R., and García-Castro, M.I. (2019). FGF Modulates the Axial Identity of Trunk hPSC-Derived Neural Crest but Not the Cranial-Trunk Decision. Stem Cell Rep. 12, 920– 933. 10.1016/j.stemcr.2019.04.015.

15. Cooper, F., Gentsch, G.E., Mitter, R., Bouissou, C., Healy, L.E., Rodriguez, A.H., Smith, J.C., and Bernardo, A.S. (2022). Rostrocaudal patterning and neural crest differentiation of human pre-neural spinal cord progenitors in vitro. Stem Cell Rep. 17, 894–910. 10.1016/j.stemcr.2022.02.018.

16. Majd, H., Amin, S., Ghazizadeh, Z., Cesiulis, A., Arroyo, E., Lankford, K., Farahvashi, S., Chemel, A.K., Okoye, M., Scantlen, M.D., et al. (2022). Deriving Schwann Cells from hPSCs Enables Disease Modeling and Drug Discovery for Diabetic Peripheral Neuropathy. 2022.08.16.504209. 10.1101/2022.08.16.504209.

17. Espinosa-Medina, I., Outin, E., Picard, C.A., Chettouh, Z., Dymecki, S., Consalez, G.G., Coppola, E., and Brunet, J.-F. (2014). Neurodevelopment. Parasympathetic ganglia derive from Schwann cell precursors. Science 345, 87–90. 10.1126/science.1253286.

18. Dyachuk, V., Furlan, A., Shahidi, M.K., Giovenco, M., Kaukua, N., Konstantinidou, C., Pachnis, V., Memic, F., Marklund, U., Müller, T., et al. (2014). Parasympathetic neurons originate from nerve-associated peripheral glial progenitors. Science 345, 82–87. 10.1126/science.1253281.

19. Adameyko, I., Lallemend, F., Aquino, J.B., Pereira, J.A., Topilko, P., Müller, T., Fritz, N., Beljajeva, A., Mochii, M., Liste, I., et al. (2009). Schwann Cell Precursors from Nerve Innervation Are a Cellular Origin of Melanocytes in Skin. Cell 139, 366–379. 10.1016/j.cell.2009.07.049.

20. Nitzan, E., Pfaltzgraff, E.R., Labosky, P.A., and Kalcheim, C. (2013). Neural crest and Schwann cell progenitor-derived melanocytes are two spatially segregated populations similarly regulated by Foxd3. Proc. Natl. Acad. Sci. U. S. A. 110, 12709–12714. 10.1073/pnas.1306287110.

21. Mica, Y., Lee, G., Chambers, S.M., Tomishima, M.J., and Studer, L. (2013). Modeling neural crest induction, melanocyte specification, and disease-related pigmentation defects in hESCs and patient-specific iPSCs. Cell Rep. 3, 1140–1152. 10.1016/j.celrep.2013.03.025.

22. Garnett, A.T., Square, T.A., and Medeiros, D.M. (2012). BMP, Wnt and FGF signals are integrated through evolutionarily conserved enhancers to achieve robust expression of Pax3 and Zic genes at the zebrafish neural plate border. Dev. Camb. Engl. 139, 4220–4231. 10.1242/dev.081497.

23. Groves, A.K., and LaBonne, C. (2014). Setting appropriate boundaries: Fate, patterning and competence at the neural plate border. Dev. Biol. 389, 2–12. 10.1016/j.ydbio.2013.11.027.

24. Jin, S., Guerrero-Juarez, C.F., Zhang, L., Chang, I., Ramos, R., Kuan, C.-H., Myung, P., Plikus, M.V., and Nie, Q. (2021). Inference and analysis of cell-cell communication using CellChat. Nat. Commun. 12, 1088. 10.1038/s41467-021-21246-9.

25. Wu, Y., Tamayo, P., and Zhang, K. (2018). Visualizing and Interpreting Single-Cell Gene Expression Datasets with Similarity Weighted Nonnegative Embedding. Cell Syst. 7, 656–666.e4. 10.1016/j.cels.2018.10.015.

26. Soldatov, R., Kaucka, M., Kastriti, M.E., Petersen, J., Chontorotzea, T., Englmaier, L., Akkuratova, N., Yang, Y., Häring, M., Dyachuk, V., et al. (2019). Spatiotemporal structure of cell fate decisions in murine neural crest. Science 364, eaas9536. 10.1126/science.aas9536.

27. Aibar, S., González-Blas, C.B., Moerman, T., Huynh-Thu, V.A., Imrichova, H., Hulselmans, G., Rambow, F., Marine, J.-C., Geurts, P., Aerts, J., et al. (2017). SCENIC: single-cell regulatory network inference and clustering. Nat. Methods 14, 1083–1086. 10.1038/nmeth.4463.

28. Kastriti, M.E., Faure, L., Von Ahsen, D., Bouderlique, T.G., Boström, J., Solovieva, T., Jackson, C., Bronner, M., Meijer, D., Hadjab, S., et al. (2022). Schwann cell precursors represent a neural crest-like state with biased multipotency. EMBO J. 41, e108780. 10.15252/embj.2021108780.

29. Buac, K., Xu, M., Cronin, J., Weeraratna, A.T., Hewitt, S.M., and Pavan, W.J. (2009). NRG1/ ERBB3 signaling in melanocyte development and melanoma: inhibition of differentiation and promotion of proliferation. Pigment Cell Melanoma Res. 22, 773–784. 10.1111/j.1755-148X.2009.00616.x.

30. Trapnell, C., Cacchiarelli, D., Grimsby, J., Pokharel, P., Li, S., Morse, M., Lennon, N.J., Livak, K.J., Mikkelsen, T.S., and Rinn, J.L. (2014). The dynamics and regulators of cell fate decisions are revealed by pseudotemporal ordering of single cells. Nat. Biotechnol. 32, 381–386. 10.1038/nbt.2859.

31. Qiu, X., Mao, Q., Tang, Y., Wang, L., Chawla, R., Pliner, H.A., and Trapnell, C. (2017). Reversed graph embedding resolves complex single-cell trajectories. Nat. Methods 14, 979–982. 10.1038/nmeth.4402.

32. Cao, J., Spielmann, M., Qiu, X., Huang, X., Ibrahim, D.M., Hill, A.J., Zhang, F., Mundlos, S., Christiansen, L., Steemers, F.J., et al. (2019). The single-cell transcriptional landscape of mammalian organogenesis. Nature 566, 496–502. 10.1038/s41586-019-0969-x.

33. Callahan, S.J., Mica, Y., and Studer, L. (2016). Feeder-free Derivation of Melanocytes from Human Pluripotent Stem Cells. J. Vis. Exp. JoVE, e53806. 10.3791/53806.

34. Solé-Boldo, L., Raddatz, G., Schütz, S., Mallm, J.-P., Rippe, K., Lonsdorf, A.S., Rodríguez-Paredes, M., and Lyko, F. (2020). Single-cell transcriptomes of the human skin reveal age-related loss of fibroblast priming. Commun. Biol. 3, 188. 10.1038/s42003-020-0922-4.

35. Rindler, K., Krausgruber, T., Thaler, F.M., Alkon, N., Bangert, C., Kurz, H., Fortelny, N., Rojahn, T.B., Jonak, C., Griss, J., et al. (2021). Spontaneously Resolved Atopic Dermatitis Shows Melanocyte and Immune Cell Activation Distinct From Healthy Control Skin. Front. Immunol. 12, 630892. 10.3389/fimmu.2021.630892.

36. Cheng, J.B., Sedgewick, A.J., Finnegan, A.I., Harirchian, P., Lee, J., Kwon, S., Fassett, M.S., Golovato, J., Gray, M., Ghadially, R., et al. (2018). Transcriptional Programming of Normal and Inflamed Human Epidermis at Single-Cell Resolution. Cell Rep. 25, 871–883. 10.1016/j.celrep.2018.09.006.

37. He, H., Suryawanshi, H., Morozov, P., Gay-Mimbrera, J., Del Duca, E., Kim, H.J., Kameyama, N., Estrada, Y., Der, E., Krueger, J.G., et al. (2020). Single-cell transcriptome analysis of human skin identifies novel fibroblast subpopulation and enrichment of immune subsets in atopic dermatitis. J. Allergy Clin. Immunol. 145, 1615–1628. 10.1016/j.jaci.2020.01.042.

38. Reynolds, G., Vegh, P., Fletcher, J., Poyner, E.F.M., Stephenson, E., Goh, I., Botting, R.A., Huang, N., Olabi, B., Dubois, A., et al. (2021). Developmental cell programs are co-opted in inflammatory skin disease. Science 371, eaba6500. 10.1126/science.aba6500.

39. Popescu, D.-M., Botting, R.A., Stephenson, E., Green, K., Webb, S., Jardine, L., Calderbank, E.F., Polanski, K., Goh, I., Efremova, M., et al. (2019). Decoding human fetal liver haematopoiesis. Nature 574, 365–371. 10.1038/s41586-019-1652-y.

40. Hughes, T.K., Wadsworth, M.H., Gierahn, T.M., Do, T., Weiss, D., Andrade, P.R., Ma, F., de Andrade Silva, B.J., Shao, S., Tsoi, L.C., et al. (2020). Second-Strand Synthesis-Based Massively Parallel scRNA-Seq Reveals Cellular States and Molecular Features of Human Inflammatory Skin Pathologies. Immunity 53, 878–894.e7. 10.1016/j.immuni.2020.09.015.

41. Kaucka, M., Szarowska, B., Kavkova, M., Kastriti, M.E., Kameneva, P., Schmidt, I., Peskova, L., Joven Araus, A., Simon, A., Kaiser, J., et al. (2021). Nerve-associated Schwann cell precursors contribute extracutaneous melanocytes to the heart, inner ear, supraorbital locations and brain meninges. Cell. Mol. Life Sci. CMLS 78, 6033–6049. 10.1007/s00018-021-03885-9.

42. Replogle, J.M., Bonnar, J.L., Pogson, A.N., Liem, C.R., Maier, N.K., Ding, Y., Russell, B.J., Wang, X., Leng, K., Guna, A., et al. (2022). Maximizing CRISPRi efficacy and accessibility with dual-sgRNA libraries and optimal effectors. eLife 11, e81856. 10.7554/eLife.81856.

43. Uhlen, M., Zhang, C., Lee, S., Sjöstedt, E., Fagerberg, L., Bidkhori, G., Benfeitas, R., Arif, M., Liu, Z., Edfors, F., et al. (2017). A pathology atlas of the human cancer transcriptome. Science 357, eaan2507. 10.1126/science.aan2507.

44. Rivero, S., Ceballos-Chávez, M., Bhattacharya, S.S., and Reyes, J.C. (2015). HMG20A is required for SNAI1-mediated epithelial to mesenchymal transition. Oncogene 34, 5264– 5276. 10.1038/onc.2014.446.

45. Lee, M., Daniels, M.J., Garnett, M.J., and Venkitaraman, A.R. (2011). A mitotic function for the high-mobility group protein HMG20b regulated by its interaction with the BRC repeats of the BRCA2 tumor suppressor. Oncogene 30, 3360–3369. 10.1038/onc.2011.55.

46. Al-Yhya, N., Khan, M.F., Almeer, R.S., Alshehri, M.M., Aldughaim, M.S., and Wadaan, M.A. (2021). Pharmacological inhibition of HDAC1/3-interacting proteins induced morphological changes, and hindered the cell proliferation and migration of hepatocellular carcinoma cells. Environ. Sci. Pollut. Res. Int. 28, 49000–49013. 10.1007/s11356-021-13668-1.

47. Savci-Heijink, C.D., Halfwerk, H., Koster, J., and Van de Vijver, M.J. (2017). Association between gene expression profile of the primary tumor and chemotherapy response of metastatic breast cancer. BMC Cancer 17, 755. 10.1186/s12885-017-3691-9.

48. Wang, T., Zhang, Y., Bai, J., Xue, Y., and Peng, Q. (2021). MMP1 and MMP9 are potential prognostic biomarkers and targets for uveal melanoma. BMC Cancer 21, 1068. 10.1186/s12885-021-08788-3.

49. Gonzalez-Avila, G., Sommer, B., Mendoza-Posada, D.A., Ramos, C., Garcia-Hernandez, A.A., and Falfan-Valencia, R. (2019). Matrix metalloproteinases participation in the metastatic process and their diagnostic and therapeutic applications in cancer. Crit. Rev. Oncol. Hematol. 137, 57–83. 10.1016/j.critrevonc.2019.02.010.

50. Gobin, E., Bagwell, K., Wagner, J., Mysona, D., Sandirasegarane, S., Smith, N., Bai, S., Sharma, A., Schleifer, R., and She, J.-X. (2019). A pan-cancer perspective of matrix metalloproteases (MMP) gene expression profile and their diagnostic/prognostic potential. BMC Cancer 19, 581. 10.1186/s12885-019-5768-0.

51. Napoli, S., Scuderi, C., Gattuso, G., Bella, V.D., Candido, S., Basile, M.S., Libra, M., and Falzone, L. (2020). Functional Roles of Matrix Metalloproteinases and Their Inhibitors in Melanoma. Cells 9, 1151. 10.3390/cells9051151.

52. Correa, B.R., de Araujo, P.R., Qiao, M., Burns, S.C., Chen, C., Schlegel, R., Agarwal, S., Galante, P.A.F., and Penalva, L.O.F. (2016). Functional genomics analyses of RNA-binding proteins reveal the splicing regulator SNRPB as an oncogenic candidate in glioblastoma. Genome Biol. 17, 125. 10.1186/s13059-016-0990-4.

53. Zhu, L., Zhang, X., and Sun, Z. (2020). SNRPB promotes cervical cancer progression through repressing p53 expression. Biomed. Pharmacother. Biomedecine Pharmacother. 125, 109948. 10.1016/j.biopha.2020.109948.

54. Liu, N., Wu, Z., Chen, A., Wang, Y., Cai, D., Zheng, J., Liu, Y., and Zhang, L. (2019). SNRPB promotes the tumorigenic potential of NSCLC in part by regulating RAB26. Cell Death Dis. 10, 667. 10.1038/s41419-019-1929-y.

55. Ma, F.-C., He, R.-Q., Lin, P., Zhong, J.-C., Ma, J., Yang, H., Hu, X.-H., and Chen, G. (2019). Profiling of prognostic alternative splicing in melanoma. Oncol. Lett. 18, 1081–1088. 10.3892/ol.2019.10453.

56. Wan, Q., Sang, X., Jin, L., and Wang, Z. (2020). Alternative Splicing Events as Indicators for the Prognosis of Uveal Melanoma. Genes 11, 227. 10.3390/genes11020227.

57. Cichorek, M., Wachulska, M., Stasiewicz, A., and Tymińska, A. (2013). Skin melanocytes: biology and development. Postepy Dermatol. Alergol. 30, 30–41. 10.5114/pdia.2013.33376.

58. Horlbeck, M.A., Gilbert, L.A., Villalta, J.E., Adamson, B., Pak, R.A., Chen, Y., Fields, A.P., Park, C.Y., Corn, J.E., Kampmann, M., et al. (2016). Compact and highly active next-generation libraries for CRISPR-mediated gene repression and activation. eLife 5, e19760. 10.7554/eLife.19760.

59. Vieth, B., Parekh, S., Ziegenhain, C., Enard, W., and Hellmann, I. (2019). A systematic evaluation of single cell RNA-seq analysis pipelines. Nat. Commun. 10, 4667. 10.1038/s41467-019-12266-7.

60. Linderman, G.C., Zhao, J., and Kluger, Y. (2018). Zero-preserving imputation of scRNA-seq data using low-rank approximation 10.1101/397588.

